# Endothelial Rap1B mediates T-cell exclusion to promote tumor growth – a novel mechanism underlying vascular immunosuppression

**DOI:** 10.1101/2021.11.03.467057

**Authors:** Guru Prasad Sharma, Ramoji Kosuru, Sribalaji Lakshmikanthan, Shikan Zheng, Yao Chen, Robert Burns, Gang Xin, Weiguo Cui, Magdalena Chrzanowska

## Abstract

Overcoming vascular immunosuppression: lack of endothelial cell (EC) responsiveness to inflammatory stimuli in the proangiogenic environment of tumors, is essential for successful cancer immunotherapy. The mechanisms through which Vascular Endothelial Growth Factor (VEGF) modulates tumor EC response to exclude T cells are not well understood. The goal was to determine the role of EC Rap1B, a small GTPase that positively regulates VEGF- angiogenesis during development, in tumor growth in vivo. Using mouse models of Rap1B deficiency, Rap1B^+/-^ and EC-specific Rap1B KO (Rap1B^i**Δ**EC^) we demonstrate that EC Rap1B restricts tumor growth and angiogenesis. More importantly, EC-specific Rap1B deletion leads to an altered tumor microenvironment with increased recruitment of leukocytes and increased activity of tumor CD8^+^ T cells. We find that tumor growth, albeit not angiogenesis, is restored in Rap1B^i**Δ**EC^ mice by depleting CD8^+^ T cells. Mechanistically, global transcriptome analysis indicated upregulation of the tumor cytokine, TNF-α, -induced signaling and NFκB transcriptional activity in Rap1B-deficient ECs. Functionally, EC Rap1B deletion led to upregulation of NFκB activity and enhanced Cell Adhesion Molecules (CAMs) expression in TNF-α stimulated ECs. Importantly, CAM expression was upregulated also in tumor ECs from Rap1B^i**Δ**EC^ mice, vs. controls. Significantly, deletion of Rap1B abrogated VEGF immunosuppressive downregulation of CAM expression, demonstrating that Rap1B is essential for VEGF-suppressive signaling. Thus, our studies identify a novel endothelial-endogenous mechanism underlying VEGF-dependent desensitization of EC to pro-inflammatory stimuli. Significantly, they identify EC Rap1 as a potential novel vascular target in cancer immunotherapy.

## Introduction

Recent cancer immunotherapies involving immune checkpoint inhibitors and chimeric antigen receptor (CAR)-T cell transfer show promise for cancer treatments. These treatments exploit the complex interactions between cancer cells and host tissue in tumor microenvironment (TME), which determine tumor growth and metastasis. Understanding the mechanisms of the host immune response to the tumor is, therefore, critical to the success of immunotherapy (Hendry et al., 2016).

Tumor vasculature is a key component of the TME. In response to hypoxia and proangiogenic cytokines, endothelial cell (EC)-driven angiogenesis gives rise to vessels that supply blood to feed the tumor. Moreover, tumor ECs control the TME by regulating leukocyte trafficking and modulating the immune response. The density of tumor-infiltrating T cells, and, in particular, CD8^+^ cytotoxic T cells, correlates with improved survival in many tumors (Fridman et al., 2012). Thus, the success of adoptive T-cell therapies to treat solid cancers depends on successful homing and infiltration by T-lymphocytes (Ager et al., 2016).

Endothelial cells control leukocyte trafficking by upregulating ligands for T-cell adhesion, among them Cell Adhesion Molecules (CAMs): VCAM-1 and ICAM-1, as a response to the proinflammatory cytokines in the TME (Ley et al., 2007). While essential for leukocyte recruitment, the proangiogenic environment in tumors dampens this response. Clinical data and experimental studies in mice suggest that tumor blood vessels are anergic to inflammatory stimuli and the recruitment of cytotoxic CD8^+^ T cells (Ager et al., 2016; Joyce and Fearon, 2015), as demonstrated in a number of tumors, including melanoma (Hendry et al., 2016).

Endothelial anergy – lack of responsiveness to inflammatory stimuli (Griffioen et al., 1996) is, in part, due to elevated angiogenic factors. In particular, VEGF-A from various TME components is upregulated in solid tumors (VEGF expression correlates with poor prognosis). VEGF decreases EC responsiveness to proinflammatory cytokines and proinflammatory adhesion receptor expression, suppressing leukocyte recruitment (Piali et al., 1995). Specifically, VEGF blocks TNF-a induced NF-kB required for CAM expression and T-cell infiltration (Huang et al., 2015). This concept is supported by evidence from anti-angiogenic therapies leading to upregulation of adhesion molecules on tumor vasculature and increased leukocyte infiltration (Dirkx et al., 2006; Hendry et al., 2016). Interference with VEGF signaling normalizes tumor vasculature and restores responsiveness of ECs to adoptive T-cell therapy also in melanoma (Shrimali et al., 2010). However, incomplete understanding of the mechanisms through which VEGF modulates tumor EC response to exclude T cells (cytokine-induced CAM expression), constitutes a major obstacle in overcoming immunosuppressive properties of tumor ECs.

Two closely related isoforms of a ubiquitously expressed small GTPase Rap1, Rap1A and Rap1B, are best known for modulation of adhesive and signaling properties of integrins and cadherins (Boettner and Van Aelst, 2009). In endothelium, Rap1 isoforms are required for vessel stabilization during development, but are not essential for vessel maintenance after birth. Instead, both isoforms are positive regulators of developmental angiogenesis (Carmona et al., 2009; Chrzanowska-Wodnicka et al., 2008; Yan et al., 2008). In particular, Rap1B, the predominant Rap1 isoform in the endothelium, promotes VEGF-induced VEGFR2 activation and signaling and its deficiency impairs VEGF-induced angiogenic responses (Chrzanowska-Wodnicka, 2010). Interestingly, Rap1B is also essential for normal VEGF-induced EC barrier dissolution and its deficiency prevents VEGF-mediated hyperpermeability in a diabetes model in vivo (Lakshmikanthan et al., 2018). Thus, there exist synergy between Rap1 and VEGF signaling. However, the role of Rap1B in tumor vasculature has not been studied and its involvement in mechanisms through which VEGF modulates tumor EC responses is unknown.

In this study our goal was to determine whether endothelial Rap1B regulated tumor growth in vivo. We hypothesized that Rap1B deficiency would lead to impaired tumor growth by restricting angiogenesis. We found that, beyond restriction of vessel growth, endothelial Rap1B deficiency led to increased leukocyte recruitment and leukocyte activation. Unexpectedly, deletion of CD8^+^ T cells rescued impaired tumor growth in EC-specific Rap1B knockout mice (Rap1B^iΔEC^), without normalizing endothelial cell number. At the molecular level, Rap1B deficiency increased EC responsiveness to TME-associated cytokine, TNFα, leading to increased T-cell adhesion. Transcriptomic analysis indicated the significant upregulation of TNFα signaling pathways and NFκB transcription, including proinflammatory CAM expression. Importantly, CAM expression was upregulated in tumor ECs from Rap1B^iΔEC^ mice. Strikingly, Rap1-deficiency prevented the VEGF-induced inhibition of ECs’ proinflammatory response. Our results demonstrate that Rap1B plays an important role in VEGF modulation of EC response in TME, acting predominantly as a suppressor of proinflammatory response. Our findings also suggest inhibition of endothelial Rap1B signaling as a novel target in overcoming endothelial anergy in tumor therapy.

## Materials and Methods

### In vivo mouse tumor model

All mouse procedures were performed according to the protocol approved by Medical College of Wisconsin Institutional Animal Use and Care Committee. Generation of total Rap1B knockout (*Rap1B*^*-/-*^) mice and endothelial cell (EC)-specific Rap1B-knockout mice (Cdh5(PAC)-CreERT2^+/0^; Rap1A^+/+^ Rap1B^f/f^; Rap1B^iΔEC^) has been previously described (Chrzanowska-Wodnicka et al., 2005; Lakshmikanthan et al., 2015). Global *Rap1B* gene heterozygous mice (*Rap1B*^*+/−*^; C57Bl/6), in which Rap1B expression is decreased to ∼50% of *Rap1B*^*+/+*^ (WT, control mice), generated by crossing male *Rap1B*^−/−^ and female WT mice used for analysis of Rap1B-deficiency on tumor growth because of high degree of lethality of total Rap1B knockout mice (*Rap1B*^*-/-*^). Cdh5-Cre-negative mice, or mice injected with carrier oil only were used as controls for Rap1B^i**Δ**EC^ mice. Both males and females between the ages of 8 and 20 weeks were used in the study. Melanoma tumors were developed with subcutaneously implanted B16F10 cells (2 × 10^5^ in 100μl PBS) into the flank area. Tumor growth was monitored using microcallipers every other day for till harvest on day 15^th^, when mice were killed, tumors isolated, weighed and processed for flow cytometry analysis. For CD8^+^-T cell depletion, tumor bearing mice were treated with purified anti-mouse CD8 mAb (clone 53-6.7), or control mAb (clone 2A3) injected itraperitoneally to mice on days 0, 2 and 9 (Figure 3A). The efficiency of CD8^+^ T cell depletion was determined in tumors, blood, bone marrow and spleen by flow cytometry and by hematological analysis.

### Cell lines and treatments

All cells lines were obtained from ATCC. B16F10 mouse melanoma cells were cultured in Roswell Park Memorial Institute (RPMI) medium supplemented with 0.1 mM nonessential amino acids, 1 mM sodium pyruvate, 2 mM l-glutamine, 25 mM HEPES, 55 μM 2-mercaptoethanol, 10% FCS and 1% penicillin-streptomycin (100 U/ml penicillin and 100 μg/ml streptomycin). Human umbilical vein endothelial cells (HUVEC) cultured in endothelial growth medium (EGM-2, Lonza, UK) for fewer than 5 passages were used in all experiments. Human T cell leukemia cell line, Jurkat E6–1, were cultured in RPMI media supplemented with 10% FCS and 1% penicillin-streptomycin. All cells were maintained in a humidified 5% CO_2_ incubator at 37 °C. 40-50% confluent HUVEC monolayers were transfected with 50nM Rap1B siGENOME siRNA pool or with non-targeting siRNA pool (Dharmacon) for 6h in OPTIMEM, and cultured for an additional 30 hours in a complete EBM culture medium. Knockdown efficiency was assessed by Western blotting. For T-cell adhesion assay, RNA sequencing and analysis of CAM expression, siRap1B or siControl-transfected HUVECs were treated with 50nM TNF-α (2% FBS EBM basal medium) for 12 hrs, prior to lysis as described below. For analysis of VEGF-induced suppression of CAM expression, cells were simultaneously treated with 50 ng/ml VEGF, as previously described (Huang et al., 2015).

### Tumor cell analysis by flow cytometry

Single-cell suspensions of tumors and peripheral organs were prepared in flow cytometry staining buffer (2% FCS + 1 mM EDTA in PBS), as follows. Tumors were digested in RPMI 1640 (Lonza) supplemented with 0.6 mg ml^−1^ collagenase type I (Worthington 4196), 0.1 mg ml^−1^ Type IV bovine pancreatic DNase (Sigma-Aldrich) and 1M MgCl_2_ (Sigma) for 45 min at 37 °C, passed through a 70 µm cell strainer and subjected to red blood cell lysis and filtration. A single-cell suspension of the spleen was obtained by tissue mincing in staining buffer. Bone Marrow (BM) cells were isolated by flushing femur and tibia bones in staining buffer and filtering through a 70μm nylon mesh. Peripheal blood monocytic cells (PBMCs) from whole blood were isolated by Histopaque gradient method. RBCs were removed using ACK lysis buffer (Lonza). The equal number of cells for each sample were stained with specific directly conjugated monoclonal antibodies (Supplemental Table 1), as previously described (Xin et al., 2020). Controls were stained with IgG isotype-matched control mAbs. 7-AAD dye (Sigma) was used to differentiate live and dead cells. Intracellular granzyme B and INF*γ* staining was performed in cells that had fixed and permeabilized in Flow Cytometry Fixation and Permeabilization Buffers (Catalog numbers FC004, FC005, R&D systems) following manufacturer’s protocol. Stained samples were analyzed on an LSR II flow cytometer (BD Biosciences) or MACSQuant Analyzer 10 (Miltenyi). The compensation was set up for all the colors used in the experiment. Data were analyzed with the FlowJo software (FlowJo LLC, version 10.0.7). Cellular debris and doublets were removed using viability dye and FSC-A FSC-H gating.

### RNA sequencing

siRap1B- or siControl-transfected HUVECs, cultured in growth media (EGM-2, Lonza) were treated with TNF-α for 12 hours. For each biological replicate RNA was extracted from 3 × 10^5^ cells grown in parallel dishes using TRIzol™ reagent. RNA quantity and quality were assessed by Agilent high sensitivity RNA-Tape Station. RNA-seq libraries were prepared using a modified SMART-Seq2 protocol (Picelli et al., 2014) and sequenced on Illumina NextSeq 500 sequencer using a high-output V2 75 cycle kit (Illumina) in 37 bp paired-end mode. Reads were pseudo-aligned to the human transcriptome (Ensembl GRCh38 release 102) and expression was quantified using Salmon v1.3.0 (Patro et al., 2017). DESeq2 v 1.26.0 Wald tests were used to determine whether gene expression fold changes were significantly different from zero (Love et al., 2014). For heatmap visualization, data were transformed using the regularized logarithmic transformation (Love et al., 2014). Pre-ranked gene set enrichment analyses (GSEA) were conducted using DESeq2 shrunken fold-changes and fgsea v1.12.0. The KEGG (Ogata et al., 1999), Reactome (Fabregat et al., 2018) and MSigDB C3:TFT (transcription factor targets) (Liberzon et al., 2011) databases were used for GSEA (Subramanian et al., 2005). The Benjamini-Hochberg method was used to adjust p-values for false-discovery in both differential expression and GSEA analysis (Benjamini and Hochberg, 1995). Data were deposited within NCBI GEO, Accession ID: GSE186046.

### NFκB activity assay

HUVECs were plated onto 6-well plates at a density of 5×10^5^ cells/well and transfected with 50 nM Rap1B siGENOME siRNA pool or with non-targeting siRNA pool (Dharmacon) for 6 hr in OPTIMEM, and cultured for an additional 18 hr in a complete EBM culture medium. After 24 hr, siControl and siRap1B cells were infected with NFκB reporter construct pNL3.2.NFκB-RE[NlucP/NF-κB-RE/Hygro] Vector (Promega) using LipofectAMINE-2000, according to the manufacturer’s protocol. pNL3.2.CMV Vector (Promega) was used as a negative control for experiments to measure regulated changes in NanoLuc^®^ luciferase expression levels. After a total of 36 hr transfection, cells were trypsinized and seeded in 96-well plates at a density of 3 or 5×10^3^ cells/well and cultured overnight in a complete EBM culture medium. Following day, cells were treated with TNF-α (20 ng/ml) or vehicle (PBS) for 5 hrs. Luciferase activity was measured using the Nano-Glo^®^ Luciferase Assay System (Promega) with an EnSpire Multimode Plate Reader (PerkinElmer), and normalized to cell number. A representative of three independent experiments, each performed in sextuplicate, is depicted as mean fold induction vs. non-stimulated cells (PBS) (Gu et al., 2016; Seigner et al., 2019).

### Adhesion assay

For quantitative assessment of T-cell adhesion to activated endothelium, we adapted previously published protocol (Wilhelmsen et al., 2013). Briefly, siRap1B or siControl-transfected HUVEC monolayers were treated with 50nM TNF-α (2% FBS EBM basal medium) for 12 hrs. Jurkat T cells were washed once with PBS and incubated in serum-free the RPMI-1640 medium with 5uM Calcein-AM (BD Biosciences-US) at 37 °C for 30 minutes. The stained Jurkat T cells were washed with PBS, and co-cultured with the TNF-α-treated HUVECs (1:1) for 45 min at 37 °C. Non-adherent Jurkat cells were removed by washing thrice with PBS. Calcein-AM fuorescence of EC-adhered Jurkat cells was immediately read on a fluorescence plate reader (Perkin Elmer EnSpire 2300) at 495/515 nm. Adhesion efficiency was calculated as a ratio of number of T cells bound per 10,000 cells plated.

### Western blotting

For determination of CAM expression, siRap1B or siControl-transfected HUVECs were treated with 50nM TNF-α (2% FBS EBM basal medium) for 12 hrs. Following completion of adhesion assay, cells were lysed in RIPA cell lysis buffer (100mM Tris-HCl buffer (pH 7.4), 0.150M NaCl, 0.001M EDTA, 1% NP-40, 1% deoxycholic acid 0.1% SDS) with 2mM sodium orthovanadate and 1X protease/ phosphatase inhibitor cocktail. Cell lysates were clarified by centrifugation and protein concentration was determined PierceTMBCA protein assay kit (Thermo Scientific). Samples containing equal amount of protein were resolved on 4-12% gradient SDS-PAGE gels and transferred to PVDF membranes. PVDF membrane was blocked in 5% skim milk/(PBS and 0.05% Tween 20) for 1 hour at RT and probed with primary antibodies against VCAM-1 (Rabbit mAb #13662) and Rap1 (Rabbit mAb #2399), both Cell Signaling Technology), and β-actin (C4: sc-47778, Santa Cruz Biotechnology) overnight at 4°C. Membranes were washed and incubated with horseradish peroxidase (HRP)-conjugated goat-anti-rabbit or donkey-anti mouse secondary antibodies for 1h at room temperature followed by chemiluminescence detection using ECL substrate (Pierce) and imaged using Amersham Imager 600 (GE Healthcare). Densitometric band intensity was determined using Amersham Imager 600 analysis software.

### Statistical analysis

Sample sizes were calculated based on variability in our and others’ previous studies. Animals were assigned to specific experimental groups without bias. When possible, subject randomization was achieved by distributing experimental groups across multiple cages and litters, as well as blinding during data collection and analysis to remove experimenter’s bias. Graphs were generated and statistical analyses performed using GraphPad Prism version 9 (GraphPad Software). Graphs were represented as mean values and standard error of mean (S.E.M.). Student’s t-test with Welch’s correction was used to measure statistical significance between groups. For comparisons of more than two groups, one-way analysis of variance (ANOVA) with Tukey’s multiple comparisons post hoc test was used. * p<0.05; ** p<0.01; *** p<0.001

## Results

### Endothelial Rap1B deletion blocks tumor growth and impairs tumor vascularization

To assess the effect of Rap1B-deficiency on tumor growth we used B16F10 melanoma model. Rap1B^+/+^ and Rap1B^+/-^ mice were each injected with 2×10^5^ B16F10 melanoma cells and tumor size was monitored until harvested for analysis on day 15 (Figure 1A). The tumor size was significantly smaller in Rap1B^+/-^ mice, starting on day 8 after inoculation (Figure 1B, D). The final tumor weight was also decreased in Rap1B^+/-^ mice compared to the wild-type control mice (Figure 1C). To determine the extent of endothelial Rap1B contribution to tumor growth, we examined melanoma growth in EC-specific Rap1B knockout (Rap1B^i**Δ**EC^) and Cdh5-Cre-negative control mice. Similar to total Rap1B depletion, endothelial deletion of Rap1B led to decreased tumor growth (Figure 1E) and decreased final weight and size of tumors (Figure 1F, G). Thus, these results suggest that the endothelial Rap1B significantly contributes to tumor growth.

**Figure 1.**
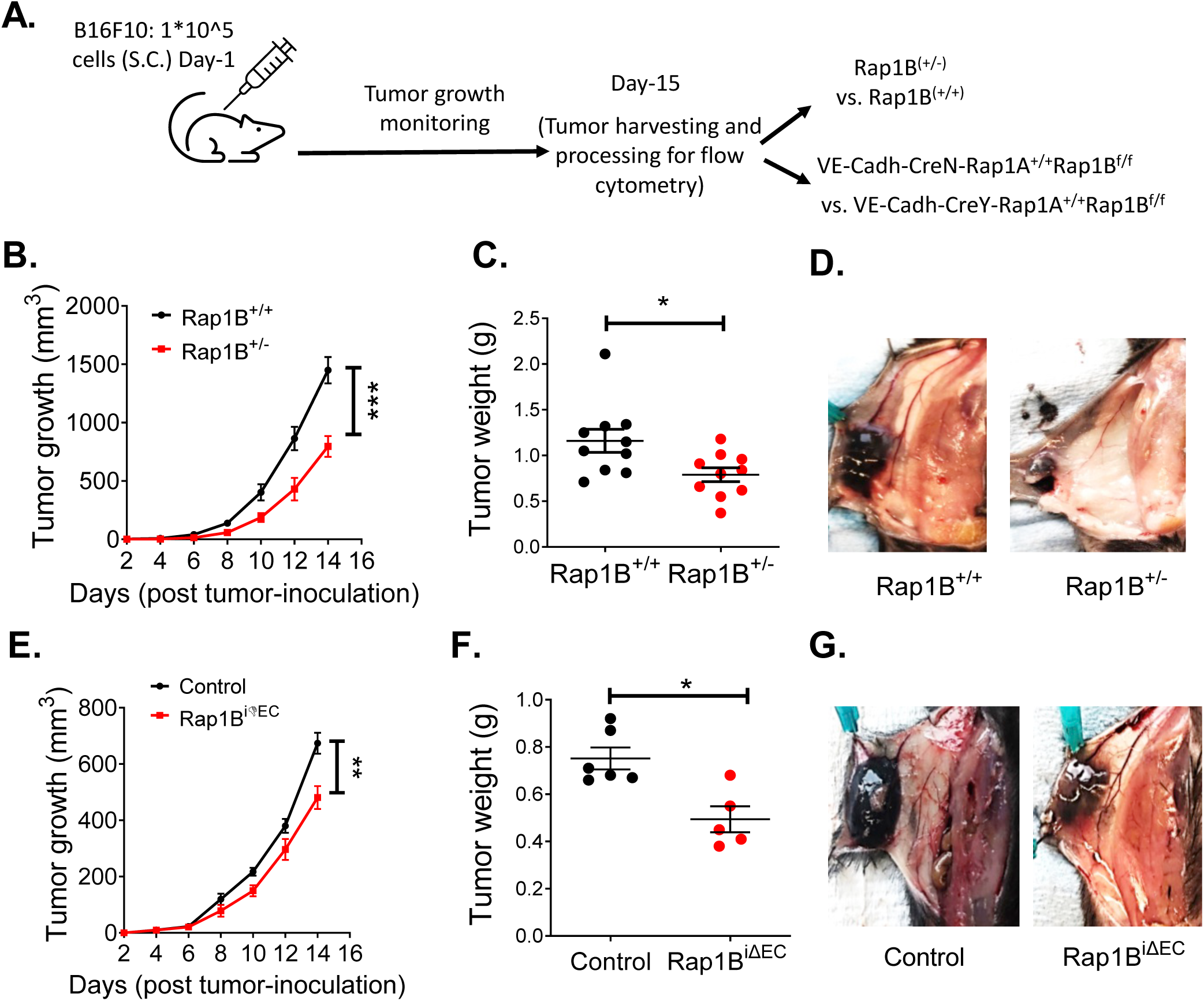
Global and EC-specific Rap1B deletion restricts tumor growth in vivo. **(A)** A schematic diagram of melanoma induction in global Rap1B-deficient mice, Rap1B^+/-^, or EC-specific Rap1B knockouts, Rap1B^iΔEC^. Tumor growth kinetics **(B)**, tumor weight at day 15 **(C)** and representative photographs of melanoma growth **(D)** in Rap1B^+/-^ (n=10) vs. Rap1B^+/+^ (n=10) mice. Tumor growth kinetics **(E)**, tumor weights **(F)** and representative photographs of melanoma growth **(G)** in Rap1B^iΔEC^ (n=5) vs. Cre-negative control mice (n=6). Data represent mean ± SEM. *, P < 0.05 **, P < 0.01, ***, P < 0.001, Student’s *t* test. Data and statistical outputs are available in *Figure 1-source data 1* file. **Figure 1-source data 1**: relates to panels A-F.

Rap1B positively regulates developmental angiogenesis by promoting EC migration and proliferation (Carmona et al., 2009; Chrzanowska-Wodnicka et al., 2008). Thus, we next examined whether EC Rap1B-deficiency impaired tumor growth by decreasing neovascularization of tumors. Tumors were excised, digested with collagenase-dispase and endothelial compartment was quantitatively assessed in CD31 (PECAM-1)-positive and CD45-negative stained cells by flow cytometry (Figure 1 – figure supplement 1A, B). We found significantly decreased number of ECs/ per tumor weights in tumors from Rap1B^i**Δ**EC^ mice, compared to control mice (Figure 2A). A similar result was obtained in tumors from Rap1B^+/-^ vs. control mice (Figure 1 – figure supplement 1C). Endothelial identity of the CD31-positive cells was further examined by immunohistochemistry in the tumor (Figure 1 – figure supplement 1D). Similar to the flow cytometry results, there was a significant decrease in CD31-positive cells in vascularized area in the tumor sections from Rap1B-deficient mice (Figure 1 – figure supplement 1E). These findings indicate that Rap1B-deficiency leads to impaired tumor endothelial cell endowment.

**Figure 2.**
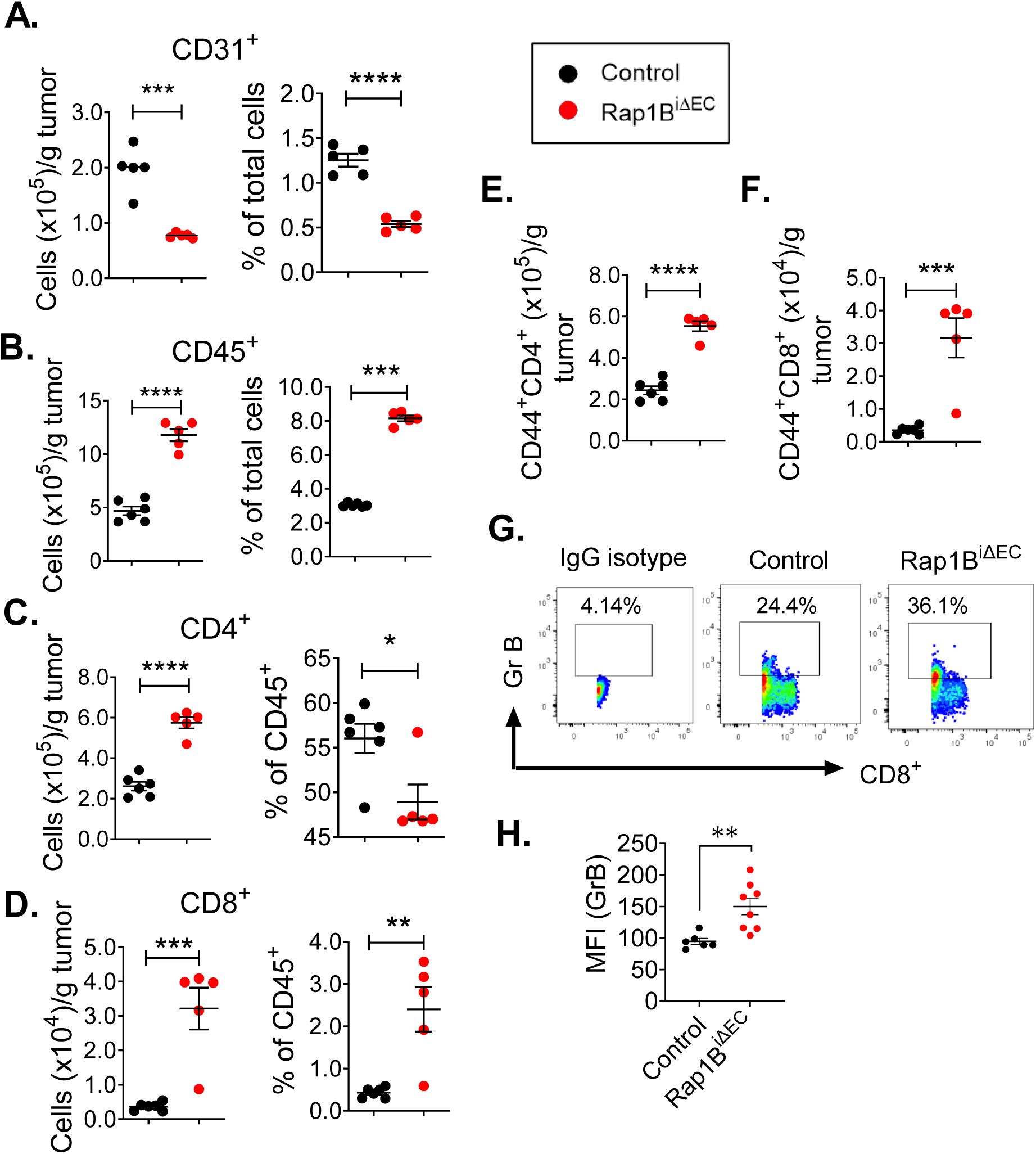
Endothelial Rap1B restricts tumor T-cell infiltration. (A-D) Endothelial Rap1B-deficiency alters TME cellularity. Flow cytometry analysis of single cell suspensions from tumors stained with cell-type specific markers demonstrate decreased endothelial cell numbers (CD31^+^, **A)**, and increased CD45^+^ cells **(B)** in Rap1B^iΔEC^ (n=5) vs. control mice (n=5). Gating strategy shown in Supplementary Fig. S1. CD4^+^ T cells **(C)** and CD8^+^ T cells **(D)** selected as shown in gating scheme in Supplementary Fig. S3. (**E-H**) **Elevated T cell activation in Rap1B^iΔEC^ vs. control mice**. Surface expression of T-cell activation marker CD44 in CD4^+^T **(E)** and CD8^+^T **(F)** cells in Rap1B^iΔEC^ (n=5) and control mice (n=6) tumors. **(G-H)** Representative FACS plots (**G**) and quantification of median fluorescence intensity, (MFI, **H**) of intracellular staining for granzyme B (GrB) in tumor CD8 ^+^ T cells from Rap1B^iΔEC^ (n=8) and control mice (n=5). Data represent mean ± SEM. *, P < 0.05 **, P < 0.01, ***, P < 0.001, Student’s t-test. Data and statistical outputs are available in *Figure 2-source data 1* file. **Figure 2-source data 1**: relates to panels A-F, H.

### Endothelial Rap1B deletion leads to increased T-cell infiltration and adhesion to Ecs

Tumor ECs-driven angiogenesis supports tumor growth by providing blood flow to the ischemic tissue. Moreover, tumor ECs actively control leukocyte infiltration and activation states. To determine if the reduced tumor growth in Rap1B^i**Δ**EC^ mice is due to impaired angiogenesis or another aspect of endothelial function altered in the absence of Rap1B, we quantitatively characterized immunce cells from TME in control and Rap1B^i**Δ**EC^ mice. Single-cell suspensions from tumors from Rap1B^i**Δ**EC^ and control mice were stained with lymphoid and myeloid cell markers and examined by flow cytometry (Figure 2 – figure supplement 1A). Interestingly, we found that EC-Rap1B deficiency led to quantitative changes in the TME cellularity (Figure 2 – figure supplement 1B-F). Specifically, the number of infiltrated CD45^+^ leukocytes (Figure 2B), and in particular, CD4^+^ (Figure 2C) and CD8^+^ (Figure 2D) T cells were increased in tumors from Rap1B^i**Δ**EC^ mice. To assess the effect of EC Rap1 deletion on CD45 activation, we examined surface expression of CD44, a cell-surface glycoprotein involved in cell-call interactions and a marker of T cell activation (Graham et al., 2007) in Rap1B^i**Δ**EC^ and control tumors. We found CD44 expression was significantly upregulated in CD4^+^ (Figure 2E) and CD8^+^ T-cells (Figure 2F) from Rap1B^i**Δ**EC^ mice. To further assess the level of T-cell activation, we examined intracellular expression of T-cell activation marker granzyme B (GrB) (Kelso et al., 2002). Consistently with increased T cell activation, we found increased number of GrB^+^ CD8^+^ T cells in tumors from Rap1B^i**Δ**EC^, vs. control mice (Figure 3G, H). Thus, EC-specific Rap1B knockout leads to decreased ECs in tumors, but increased leukocyte infiltration and increased leukocyte activity.

**Figure 3.**
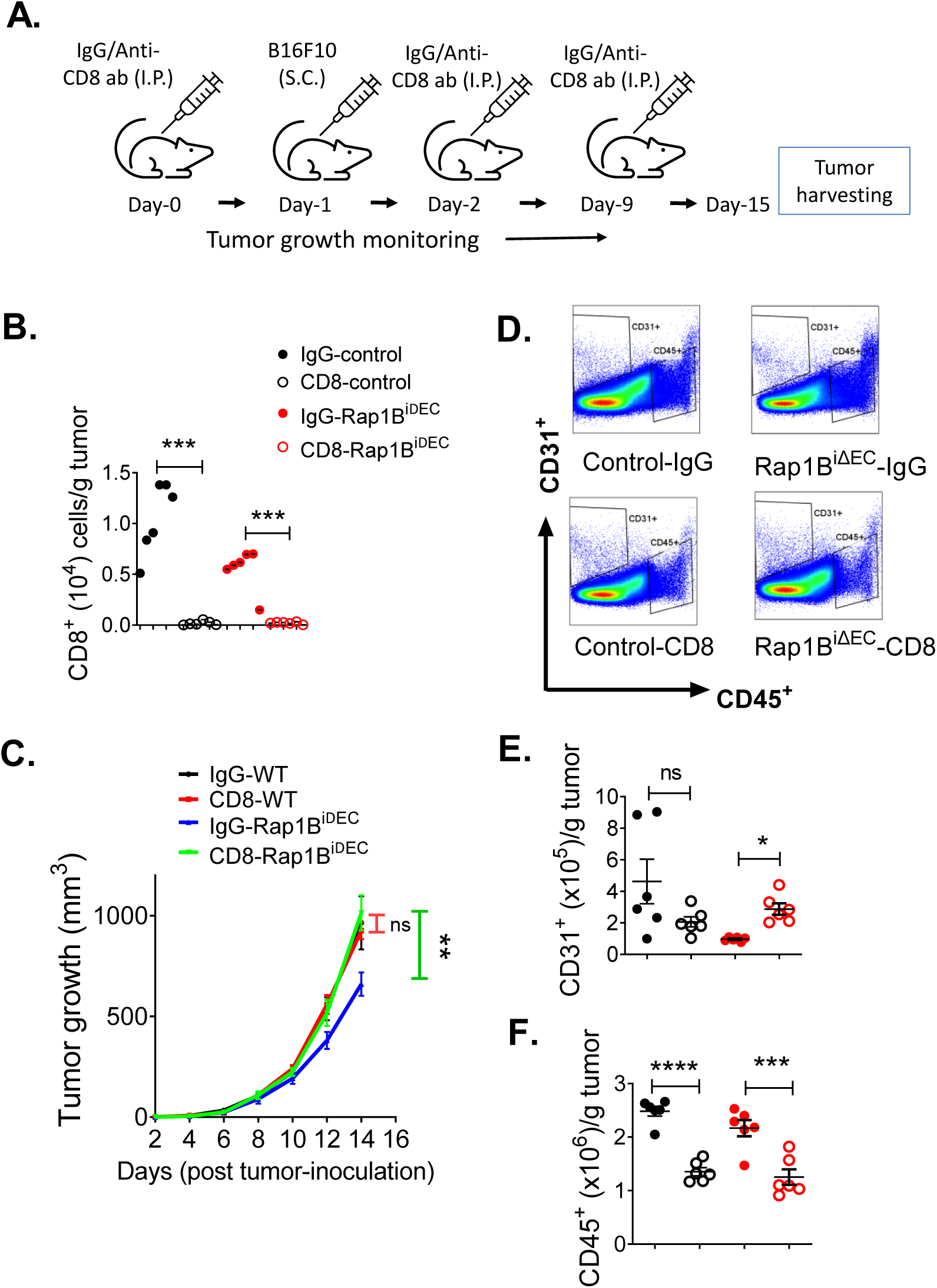
Depletion of CD8^+^ T cells normalizes tumor growth in Rap1B^iΔEC^ mice. Schematic diagram of antibody treatment (intraperitoneal, I.P. injection) and melanoma induction (subcutaneous,S.C. injection). **(B)** Quantification of CD8^+^ T cells demonstrates highly efficient cell depletion in with anti-CD8 antibody or IgG isotype control. **(C)** Tumor growth kinetics in Rap1B^iΔEC^ vs. Cre-negative control treated with anti-CD8. **(D)** Gating scheme and quantification of CD31^+^ ECs **(E)** and CD45^+^ cells **(F)**. Data are presented as the mean ± S.E.M (n=6 per group). *, P < 0.05, **, P < 0.01, ***, P < 0.001, Panels B, C, F: one-way ANOVA with Tukey’s post hoc test. Panel E: Student’s t-test. Data and statistical outputs are available in *Figure 3-source data 1* file. **Figure 3-source data 1**: relates to panels B, C, E and F.

### Depletion of CD8^+^ T cells rescues tumor growth in Rap1B^iΔEC^ mice

Increased presence of CD8^+^ cytotoxic T cells is associated with reduced tumor growth (Ager et al., 2016; Joyce and Fearon, 2015). Our results showing increased leukocyte activation and recruitment to tumors in Rap1B^i**Δ**EC^ mice suggested endothelial Rap1B might control tumor growth by restricting CD8^+^ T cell recruitment and activation. To determine if increased CD8^+^-T-cell activity/infiltration was responsible for suppressed tumor growth in Rap1B^i**Δ**EC^ mice, we examined the effect of CD8^+^ T-cell depletion *in vivo* on tumor growth and TME cellularity in Rap1B^i**Δ**EC^ and control mice (Figure 3A). Injection of anti-CD8*α* monoclonal antibody (mAb), successfully depleted CD8^+^ T cells (Figure 3B), but not CD4^+^ T cells (Figure 3 – figure supplement 1A) in tumors from control and Rap1B^i**Δ**EC^ mice. CD8^+^ T cells were also depleted in other tissue compartments (blood, spleen and bone marrow, Figure 3 – figure supplement 2). Consistent with T-cell depletion, the overall number of CD45^+^ cells was also decreased (Fig. 3F, Figure 3– figure supplement 3). Interestingly, we found that the anti-CD8*α* Ab treatment did not lead to decreased tumor growth in control mice, compared to mice treated with isotype control mAb (Figure 3C). This further supports the view that in endothelial Rap1 restricts CD8^+^ T cells’ effector function and tissue homing in this aggressive melanoma mouse model. In contrast, anti-CD8*α* mAb injection restored tumor growth in Rap1B^i**Δ**EC^ mice to the level of control mice (Fig. 3C, Figure 3 – figure supplement 4).

We considered a possibility that restored tumor growth in Rap1B^iΔEC^ mice upon depletion of CD8^+^ T cell may be via increased angiogenesis (increased EC endowment) rather than altering of CD8^+^ T cell -EC interactions. To this end, we examined EC endowment in tumors from control and Rap1B^i**Δ**EC^ mice (Figure 3D). Although CD8^+^ T-cell depletion did not retard tumor growth in normal mice, it resulted in a significant decrease in the number of tumor ECs, compared to isotype control mAb-injected mice (Figure 3E). This suggests that in control tumors, EC number is not a tumor growth-limiting factor. Interestingly, depletion of CD8^+^ T cells lead to a small, but significant elevation of ECs numbers in tumors in Rap1B^i**Δ**EC^ mice (Figure 3E), albeit to a level lower than in WT IgG controls. These results show that restored tumor growth in Rap1B^i**Δ**EC^ mice is not likely due to improved angiogenesis, but rather inhibition of increased CD8^+^ T cell activity upon deletion of Rap1B from endothelium.

### Rap1B suppresses TNFα-induced NFκB transcription and signaling

To gain insights into the mechanism through which Rap1B regulates the observed increased leukocyte recruitment and activity, we analyzed the global effects of the TME-associated cytokine, TNF-α treatment on gene expression in siRap1B and siControl ECs by bulk RNA sequencing. Principle component and differential gene expression analyses showed that siRap1B ECs exhibited a distinct transcriptional profile compared to siControl ECs (Figure 4 – figure supplement 1A, B). To assess the TNF-α-induced changes in ECs upon Rap1B knockdown, we conducted gene set enrichment analysis (GSEA) using gene sets from the Reactome database. This analysis revealed upregulation of genes in the TNF-α signaling pathway in siRap1B ECs (Figure 4A, B). Furthermore, the transcriptional profile was consistent with upregulation of NFκB transcriptional activity in siRap1B cells (Figure 4C, D and Figure 4 – figure supplement 1C, D). To examine the effect of Rap1B deletion on NFκB activity in ECs, siControl and siRap1B HUVECs were transfected with a NFκB-driven luciferase reporter plasmid or a control, empty plasmid, and emitted luminescence was measured after 5h cell exposure to TNF-α (20 ng/ml). TNF-α induced a modest, but significant luminescence increase in siControl ECs, vs. PBS (Figure 4E). Strikingly, TNF-α induced luminescence was significantly higher in siRap1B ECs (Figure 4E). Consistently with increased NFκB activity, GSEA revealed upregulation of NFκB targets (Figure 4F).

**Figure 4.**
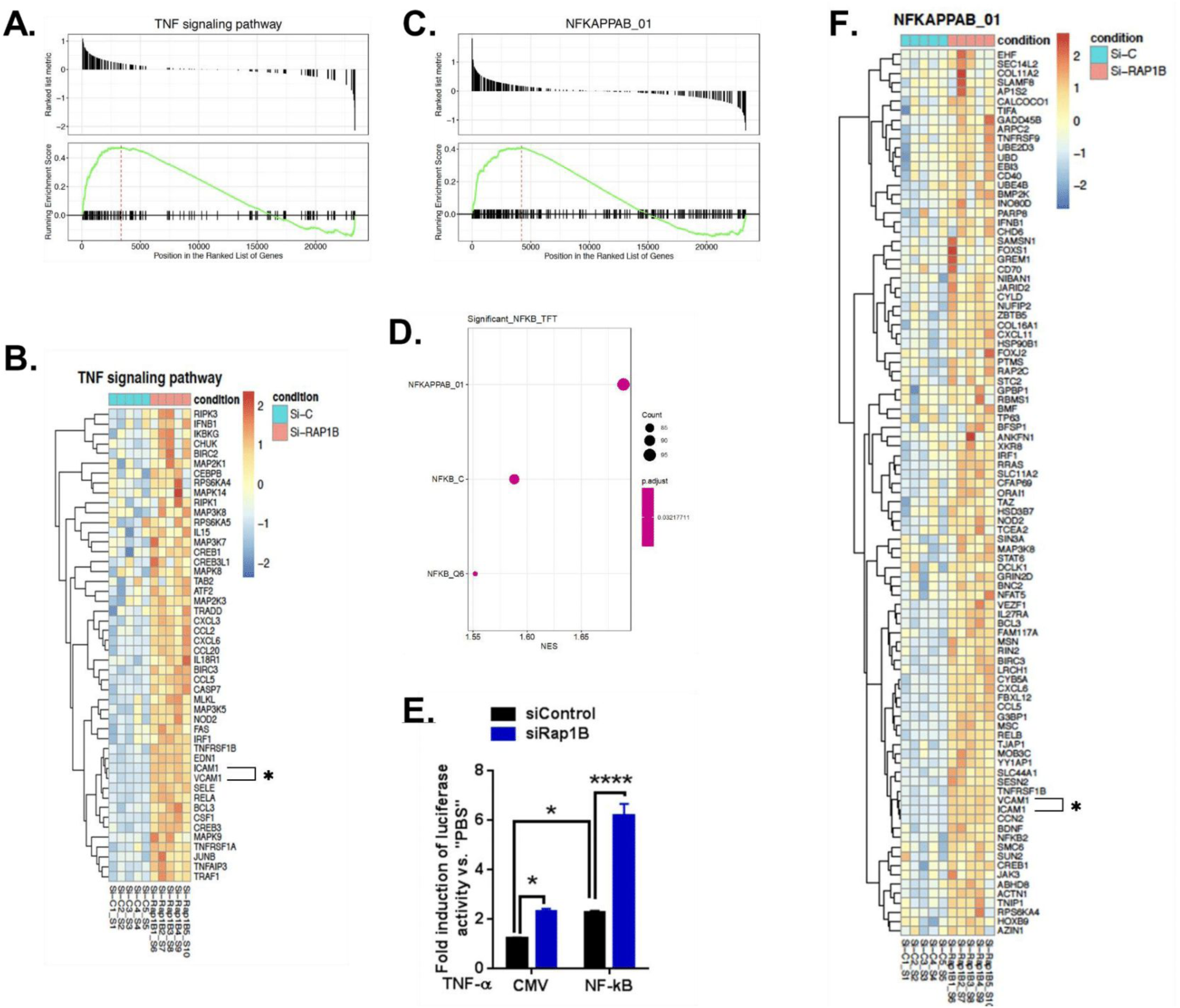
Endothelial Rap1B restricts TNF-α induced NFκB activation. Increased TNF-α signaling and NFκB transcription detected by bulk RNA sequencing of TNF-α-treated siRap1B and siControl Ecs. **(A)** Gene set enrichment analysis (GSEA) of genes in TNF-α signaling pathway upregulated in siRap1B vs. siControl. **(B)** Heatmap of selected genes from TNF-α signaling pathway. **(C)** GSEA of NFκB signaling pathway. **(D)** Transcriptional activation of NFκB in siRap1B ECs. **(E)** TNF-α-induced NFκB activity determined by luciferase assay in cells transfected with NF-κB reporter construct or a control vector (CMV). Values shown as mean fold change vs. PBS-treated cells. Error bars represent mean ± SD (n = 6). *, P < 0.05, ****, P < 0.0005. One-way ANOVA followed by Tukey’s multiple comparisons test. **(F)** Heatmap of selected genes from NFκB pathway. (n = 5 separate EC sets per group). Link to sequence data deposited within NCBI GEO is available in *Figure 4-source data 1* file. Data and statistical outputs for panel E are available in *Figure 4-source data 2* file. **Figure 4-source data 1**: relates to panels A-D and F. **Figure 4-source data 2**: relates to panel E.

Notably, among NFκB-regulated genes upregulated in siRap1B ECs were proinflammatory Cell Adhesion Molecules (CAM)s; ICAM-1 and VCAM-1 (Figures 4B and F, asterisks). To validate these RNAseq data, we examined the effect of TNF-α on CAM protein expression in siRap1B and siControl ECs. TNF-α treatment led to a significant upregulation of VCAM-1 (Figure 5A) and ICAM-1 (Figure 5– figure supplement 1A) in control ECs. Interestingly, and consistent with increased transcript expression, TNF-α-induced CAM protein expression was significantly enhanced in Rap1B-deficient ECs (Figure 5A, Figure 5– figure supplement 1A). These findings show that in isolated ECs deletion of Rap1B leads to upregulation of CAMs via upregulation of NFkB signaling in response to TME cytokine, TNF-α. Importantly, we found that, similarly to the *in vitro* effect, EC-Rap1B deletion mice led to elevated expression of VCAM (Figure 5B, C) and ICAM-1 (Figure 5– figure supplement 1B) in tumor ECs from Rap1B^iΔEC^ vs. control mice. Thus, both *in vitro* and *in vivo*, EC Rap1B restricts CAM expression in response to a TME-associated cytokine.

**Figure 5.**
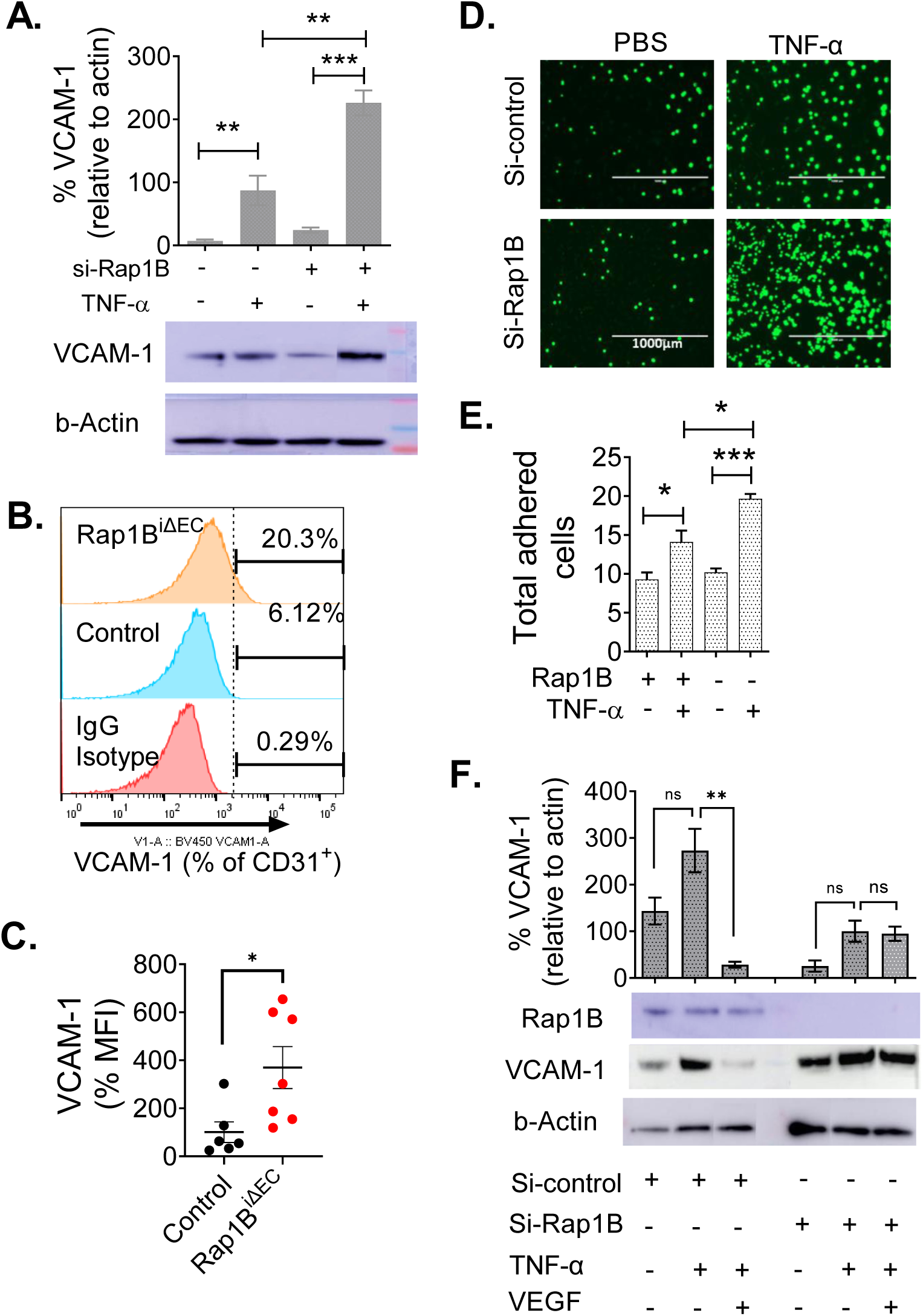
Endothelial Rap1B restricts proinflammatory CAM expression, mediates VEGF signaling. **(A)** Top: Densitometry of TNFα-induced VCAM-1 expression, normalized to actin, in siControl- or siRap1B-transfected HUVECs. Bottom: representative Western blot. Original images are available in *Figure 5-source data 1* file. **(B, C)** VCAM-1 expression in tumor ECs (CD31^+^ cells): representative flow cytometry histogram (**B**) and median fluorescence intensity (MFI) (**C**). n ≥ 5 mice per group, **(D, E)** EC-Rap1B deficiency leads to increased leukocyte adhesion in vitro. Representative image (**D**) and quantification (**E**) of Calcein-AM-labeled Jurkat cells adhering to siRap1B and siControl ECs following TNF-α treatment 12 hours. Adhesion efficiency is expressed as Jurkat cells bound/10,000 plated. (**F**) VEGF treatment inhibits TNF-α-induced CAM expression in siControl but not in siRap1B ECs. Densitometry (top) and representative Western blot (bottom) are shown. Original images are available in *Figure 5-source data 2* file. Data shown are mean ± S.E.M. *, P < 0.05, **, P < 0.01, ***, P < 0.001, Panels A, E, F: one-way ANOVA followed by Tukey’s multiple comparisons test. Panel C: Student’s t-test. Data and statistical outputs are available in *Figure 5-source data 3* file. **Figure 5-source data 1**: original images of Western blots in panel A. **Figure 5-source data 2**: original images of Western blots in panel F. **Figure 5-source data 3:** relates to panels A, C, E and F.

EC-controlled expression of CAMs on EC surface controls leukocyte infiltration into tumors, by promoting leukocyte adhesion via integrins on leukocytes and CAMs on endothelium. To determine if increased adhesion to Rap1B-deficient endothelium may contribute to increased tumor infiltration, we measured the effect of TNF-α on leukocyte adhesion in Rap1B-deficient and WT ECs (Wilhelmsen et al., 2013) (Figure 5D). A 24-hour treatment with TNF-α led to a significant increase in leukocyte adhesion in control cells (Figure 5E). However, the number of leukocytes adhering to ECs was significantly increased upon Rap1B knockdown (Figure 5E), supporting our hypothesis that deletion of Rap1B in EC promotes EC-leukocyte interactions. In sum, underlying increased T-cell infiltration into Rap1B^iΔEC^ tumors is elevated EC inflammatory NFkB activation and CAM expression, which leads to increased adhesion and activation of leukocytes.

### Endothelial Rap1B mediates immune suppression in the VEGF pathway

In addition to stimulating EC angiogenic responses, VEGF promotes tumor growth by dampening EC responsiveness to proinflammatory cytokines and proinflammatory adhesion receptor expression, suppressing leukocyte recruitment. Because Rap1B is a positive regulator of VEGF-mediated angiogenic responses (Chrzanowska-Wodnicka, 2010; Lakshmikanthan et al., 2011), we hypothesized Rap1B may mediate VEGF-induced suppression of EC pro-inflammatory responses. To test this hypothesis, we examined the effect of Rap1B knockdown on VEGF-mediated modulation of VCAM-1 expression. Consistent with previous reports (Huang et al., 2015), TNF-α-induced VCAM-1 expression was largely inhibited in the presence of VEGF (Fig. 5F). Strikingly, this inhibitory effect of VEGF was abolished in siRap1B ECs, where VEGF failed to reduce TNF-α-induced VCAM-1 expression (Figure 5F). Therefore, Rap1B is an essential component of the VEGF signaling that modulates EC response to proinflammatory cytokines. Our findings suggest Rap1B may be a key EC signaling component mediating tumor EC anergy (vasosuppression).

## Discussion

Overcoming vascular immunosuppression - reduced EC responsiveness to inflammatory stimuli - is essential for successful cancer immunotherapy. In this manuscript we demonstrate Rap1B is a key EC component restricting endothelial proinflammatory response. We show that EC Rap1B is permissive for tumor growth, as its EC deficiency suppresses tumor growth and angiogenesis. Perhaps more importantly, the deletion of Rap1B leads to altered TME, increased leukocyte recruitment, and, in particular, increased activity of tumor CD8^+^ T cells. We find that tumor growth, albeit not angiogenesis, is restored in Rap1B^i**Δ**EC^ mice by depleting CD8^+^ T cells. Therefore, the key function of Rap1B in tumor ECs is to control leukocyte, and specifically CD8^+^ T cell interactions. Mechanistically, we show that Rap1B deletion leads to upregulation of the TNF-α signaling and cytokine-induced NFκB transcriptional activity. Specifically, EC Rap1B restricts proinflammatory cytokine-induced upregulation of CAMs on ECs and EC-leukocyte interactions *in vitro* and in tumor ECs, *in vivo*. Importantly, in the proangiogenic environment present in tumors, Rap1B is essential for mediating VEGF-immunosuppressive signaling, as Rap1-deficiency inhibits VEGF-mediated inhibition of CAM expression. Thus, our studies identify a novel EC-endogenous mechanism underlying VEGF-dependent desensitization of ECs to proinflammatory stimuli. Significantly, they identify EC Rap1B as a potential novel vascular adoptive cell therapy target.

Increased angiogenic factors in TME are linked with tumor progression and tumor vasculature has been a target of anti-tumor therapies (Ferrara and Adamis, 2016; Hendry et al., 2016). Anti-angiogenic therapies block vessel growth, normalize vascular permeability and promote the recruitment of pericytes to stabilize blood flow. Importantly, they sensitize anergic tumor endothelial cells to proinflammatory stimuli, induce proinflammatory expression program which promotes tumor-infiltrating T cells, a positive are prognostic for patient outcome in multiple cancers (Fridman et al., 2012). VEGF is the most prominent tumor angiogenic factor and a number of anti-VEGF therapies have been approved for treatment of cancer (Joyce and Fearon, 2015). However, anti-VEGF therapies lead to adverse effects (tumor aggressiveness, increased metastasis (Ebos et al., 2009; Pàez-Ribes et al., 2009). Moreover, while molecular pathways underlying VEGF-induced vessel formation have been intensely investigated, the mechanism through which VEGF modulates immune responses of tumor vessels remains poorly understood.

The findings described in this manuscript provide novel molecular insights into the molecular underpinnings of VEGF-induced immunosuppression. Previous studies implicated Rap1B developmental angiogenesis (Chrzanowska-Wodnicka, 2013) and demonstrated that Rap1B controls multiple aspects of EC angiogenic responses to VEGF (Chrzanowska-Wodnicka, 2010). More recent, molecular studies demonstrated Rap1 acts as an upstream, positive regulator of VEGFR2 signaling important for shear stress sensing and NO release from ECs (Lakshmikanthan et al., 2015), and VEGF-induced vascular permeability (Lakshmikanthan et al., 2018). Consistently with that model, EC-Rap1B deficiency attenuates VEGF-induced permeability *in vivo* and protects from VEGF-induced hyperpermeability in a diabetes model (Lakshmikanthan et al., 2018). Here, we demonstrate, for the first time, to our knowledge, that Rap1 is essential for VEGF-induced inhibition of the proinflammatory response. While the exact molecular mechanisms need to be elucidated, our findings place Rap1B at the crossroads of VEGFR2 and cytokine signaling, essential for control of tumor growth (Fig. 6).

**Figure 6.**
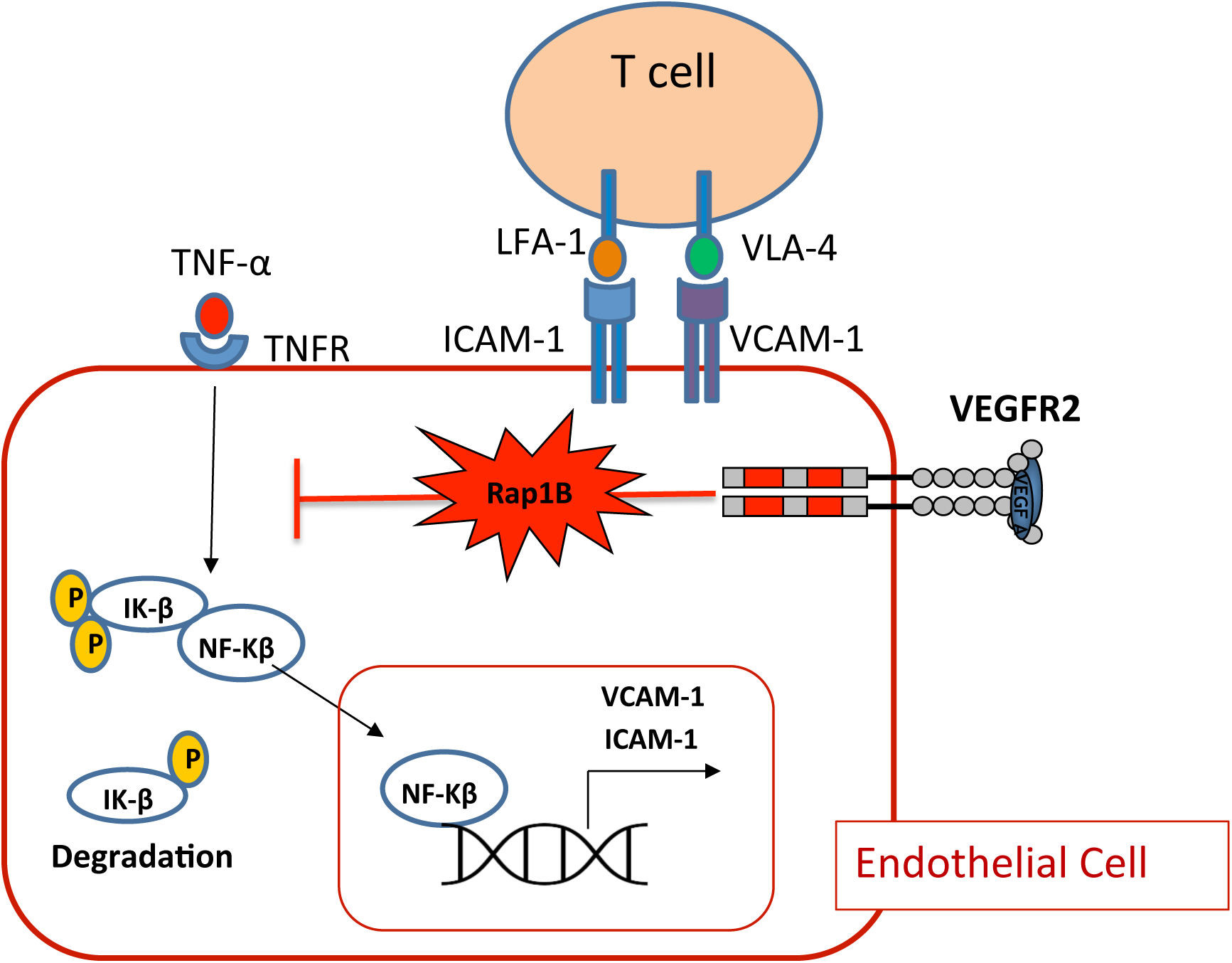
Endothelial Rap1B conveys VEGF suppression of immune reactivity – proposed mechanism. Under normal conditions Rap1 suppresses cytokine-induced CAM expression, limiting T-cell adhesion and recruitment. In proangiogenic conditions of TME, VEGF signaling, mediated by Rap1B, suppresses this endothelial immune response.

An unbiased transcriptomics approach provided a novel, global view of pathways controlled by Rap1B (Figure 4 and Figure 4 – figure supplement 1). The top pathway upregulated in the absence of Rap1B is the NFκB -mediated proinflammatory response. Deletion of Rap1B leads to upregulation of NFκB targets, including CAM receptors, both: *in vitro* and in tumor ECs *in vivo*. Interestingly, Rap1B deletion also leads to upregulation of CAMs in a mouse atherosclerosis model (Singh et al., 2021). Additional NFκB proinflammatory targets likely contribute to the increased leukocyte infiltration and leukocyte infiltration in Rap1B^i**Δ**EC^ tumors, and warrant further investigation. The analysis of Rap1B^i**Δ**EC^ TME cellularity revealed that EC Rap1B-deficiency selectively affected other leukocyte populations; in particular the number of some tumor B cells was increased in Rap1B^i**Δ**EC^ tumors vs. controls (Figure 2 – figure supplement 1D, E). Future studies will determine whether these effects are primary or secondary. Nonetheless, our findings support Rap1B as a key regulator of EC-leukocyte interactions via a novel mechanism involving suppression of proinflammatory signaling. These results also reveal a new aspect of Rap1 biology in endothelium. Future studies will focus on the mechanisms of transcriptional control by Rap1B and their implications in proinflammatory states.

In sum, we identified Rap1B as a novel molecular regulator of EC proinflammatory response, which may lead to new therapeutic approaches to overcome vascular immunosuppression in cancer immunotherapy.

## Figure, figure supplements and figure source data legends

**Figure 1 – figure supplement 1.**
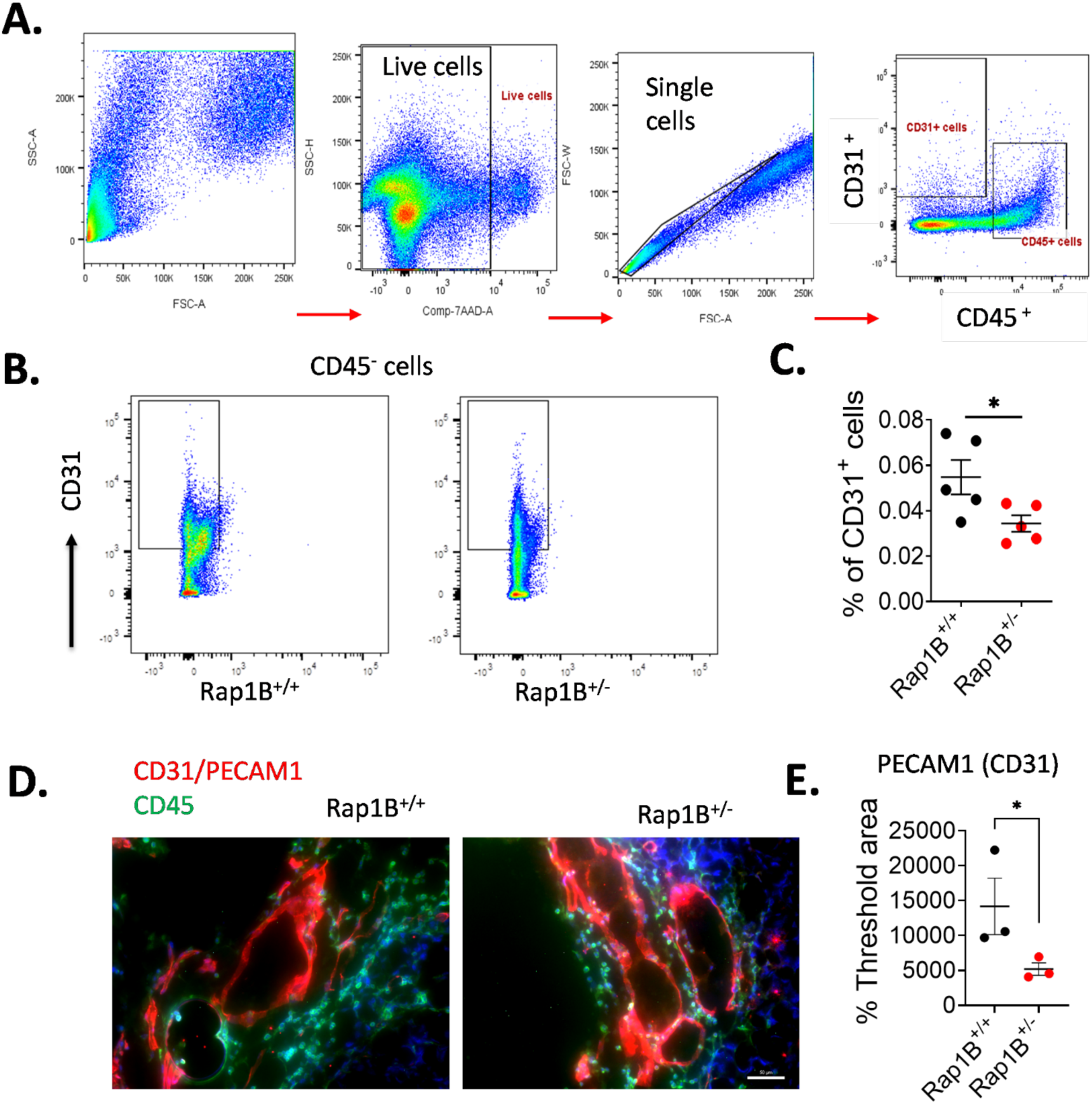
Reduced CD31^+^ endothelial cells in Rap1B^+/-^ tumors. (**A**). Gating strategy for endothelial cells (CD31^+^) and leukocytes (CD45^+^) using stained single-cell suspension from the harvested tumors. Representative plots from a control tumor are shown. One million events were acquired on BD FACS-diva (LSR II-Green) using single-cell suspension from the harvested tumors. Viable, 7-AAD-negative cells were identified, and cell doublets were discriminated by their SSC and FSC characteristics. Single cells were further gated for leukocytes (CD45^+^/CD31^-^) and endothelial cells (CD31^+^/CD45^-^). (**B**) Decreased number of tumor endothelial cells (CD31^+^/CD45^-^) isolated from Rap1B^+/-^ mice vs. control, shown as % of gated single cells (**C** *, P < 0.05). Data are presented as the mean ± S.E.M. n = 5 mice per group, student’s *t* test. (**D**) Typical immunofluorescent staining of CD45^+^ cells (green) and CD31^+^ endothelial cells (red) in tumor sections. (**E**) CD31^+^ endothelial cells quantification demonstrates a significant decrease in ECs in tumor sections in Rap1B^+/-^, vs. control group (*, P < 0.05). n = 3 mice per group, Student’s *t* test. Data and statistical outputs are available in *Figure 1 – figure supplement 1-source data 1* file. **Figure 1 – figure supplement 1-source data 1:** relates to panels C and E.

**Figure 2-figure supplement 1.**
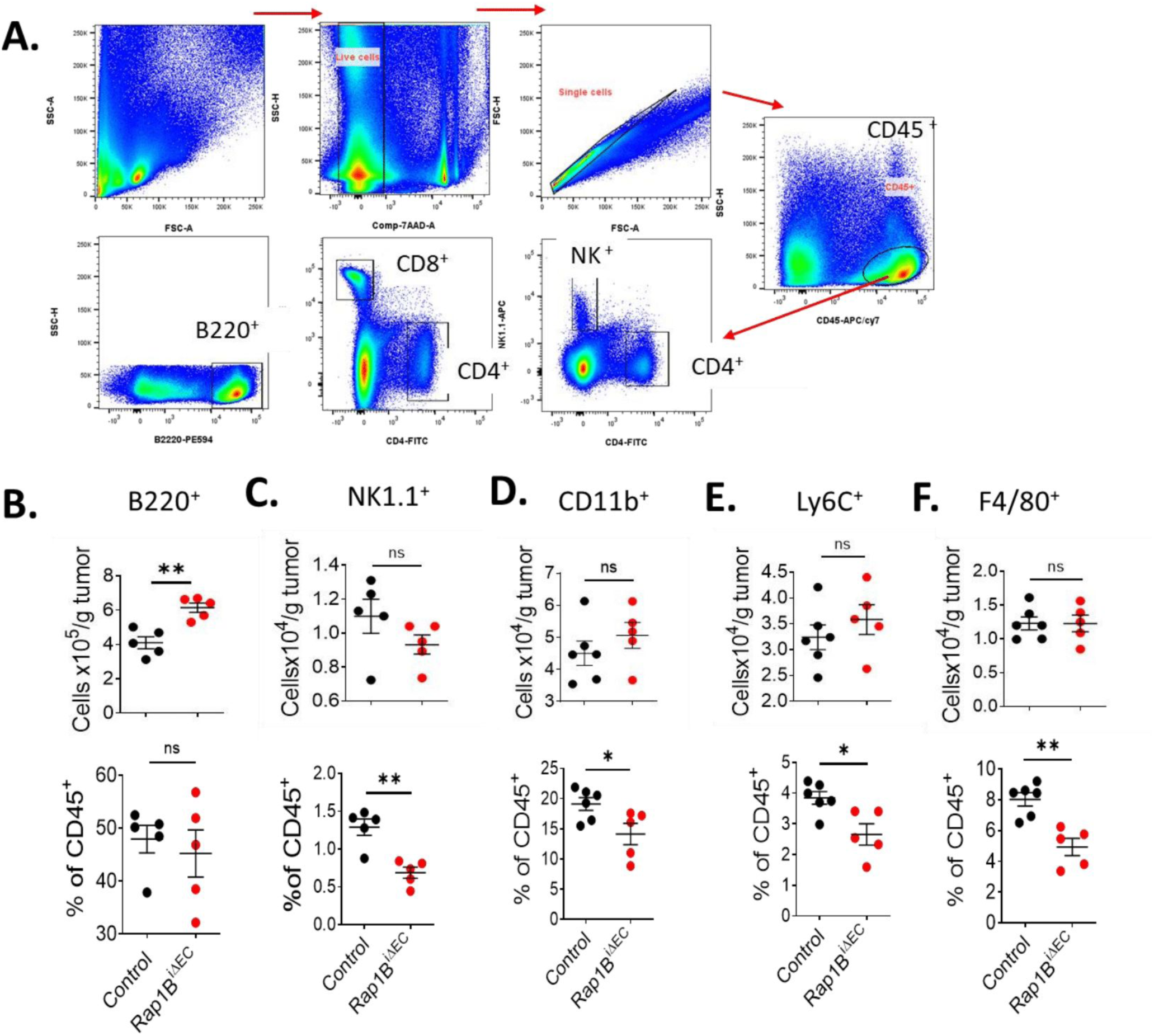
EC-specific Rap1B deletion alters recruitment of tumor-infiltra:ng lymphocytes (TIL). (**A**) Gating and selection of immune cell types from tumor single-cell suspensions. Representative plots from a control tumor are shown. One million events were acquired on BD FACS-diva (LSR II-Green) using single-cell suspension from the harvested tumors. Single cells were used to identify CD45^+^ leukocytes, which were further gated using marker-specific antibodies against NK1.1 (natural killer cells), CD4, CD8 and B220 B-cells. **B-F**. Quantitation of cell populations: **(B)** B cells (B220^+^), **(C)** natural killer cells (NK1.1^+^), **(D)** myeloid cells (CD11b^+^), **(E)** monocytes (Ly6C^+^) and macrophage (F4/80^+^) in single-cell suspension of tumors from Rap1B^iΔEC^ or control mice. Data are presented as the mean ± S.E.M. *, P < 0.05. n = 5 mice per group, Student’s *t* test. Data and statistical outputs are available in *Figure 2 – figure supplement 1-source data 1* file. **Figure 2– figure supplement 1-source data 1**: relates to panels B-F.

**Figure 3 – figure supplement 1.**
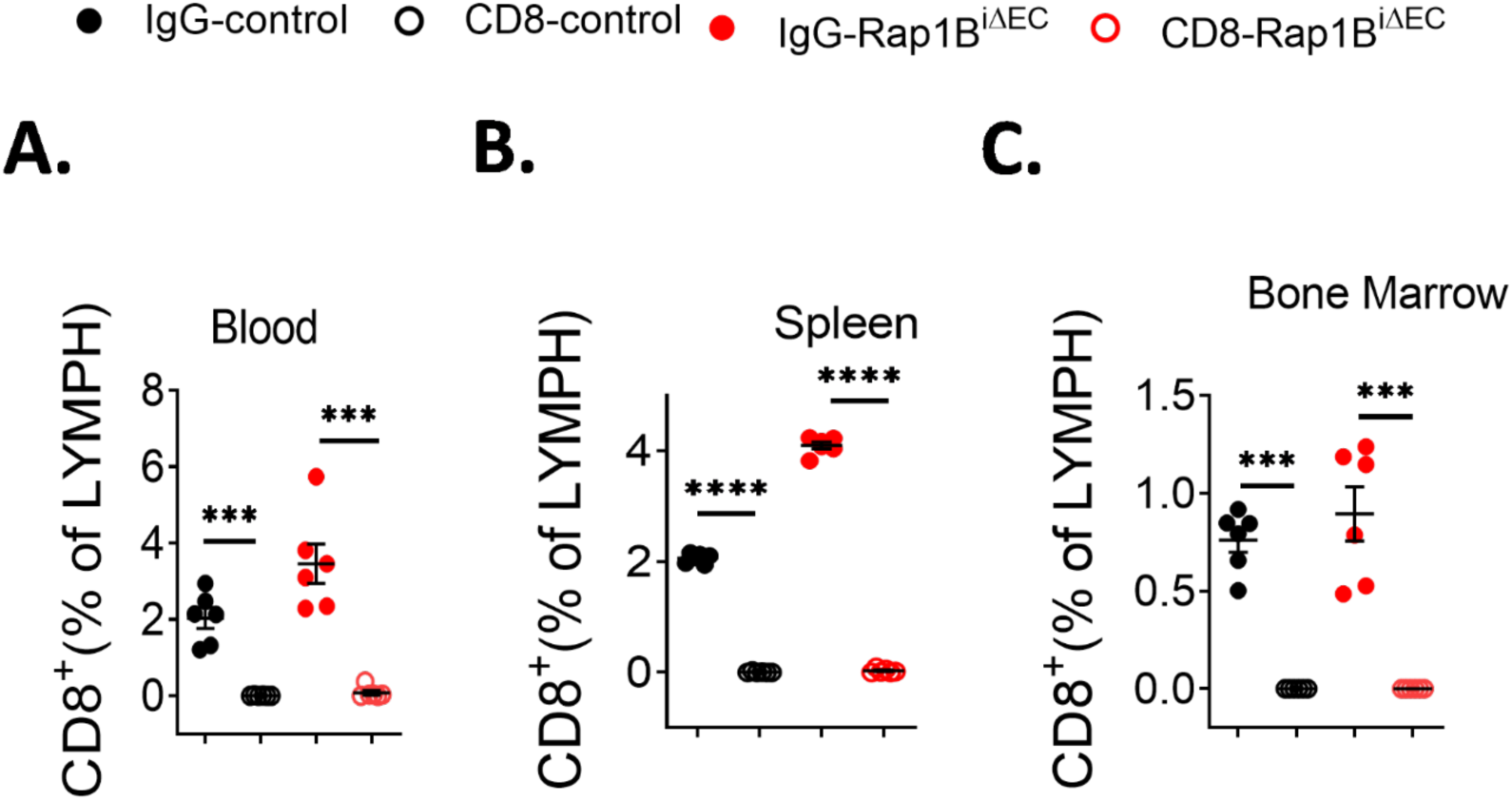
Efficiency of CD8^+^ T cell depletion in specific :ssue compartments. Peripheral blood cells **(A)**, spleen cells **(B)** and bone marrow cells (**C)** were stained with Very CD8 mAb. Flow analysis showing drastic depletion of CD8^+^ cells in all tissue compartment. IgG isotype control has no significant modulation on T cell. Data are presented as the mean ± S.E.M. ***, P < 0.001, ****, P < 0.0001. n = 6 mice per group, one-way ANOVA with Tukey’s multiple comparisons post hoc test. Data and statistical outputs are available in *Figure 3 – figure supplement 1-source data 1* file. **Figure 3 – figure supplement 1-source data 1:** relates to panels A-F.

**Figure 3 – figure supplement 2.**
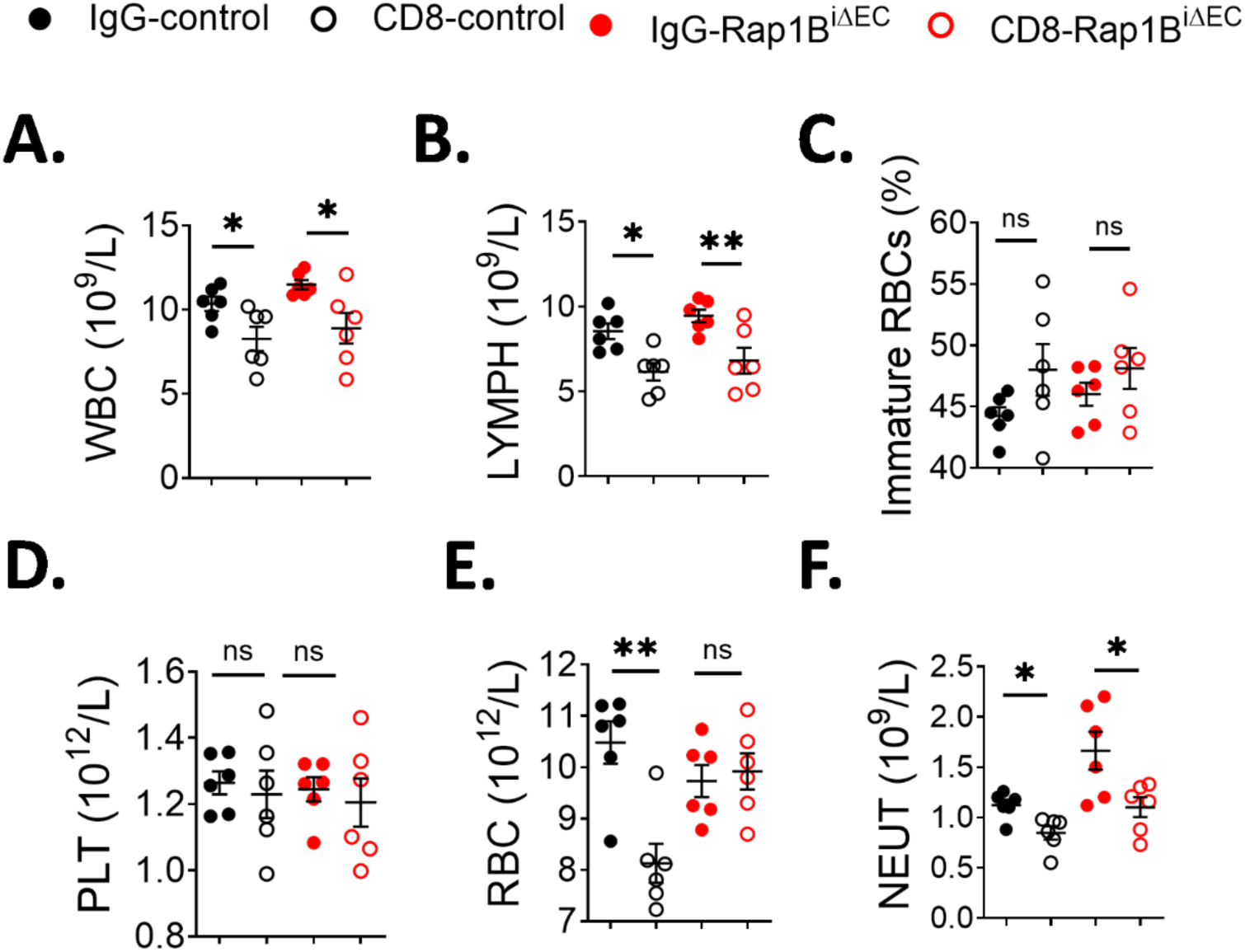
Hematological (CBC) analysis in control and Rap1B^iΔEC^ mice aTer CD8^+^ T cell depletion. Whole blood (13 µl per mouse) was collected into anticoagulant (1:10) using capillary tubes (StatSpin^®^ Microhematocrit Tubes, VWR). The blood samples were immediately analyzed using an automated Hematology blood analyzer (Sysmex XN-1000™). CBC counts: **(A)** White blood cells (WBC) 10^9^/L; **(B)** lymphocyte (LYMPH) 10^9^/L. **(C)** Immature (young) RBC fraction, as measured by % of high fluorescence reticulocyte (HFR), **(D)** Platelets (PLT) 10^12^/L, **(E)** Red blood cells (RBC) 10^12^/L, and **(F)** Neutrophils (NEUT). Data are presented as the mean ± S.E.M. *, P < 0.05, **, P < 0.01. n = 6 mice per group, one-way ANOVA with Tukey’s multiple comparisons post hoc test. Data and statistical outputs are available in *Figure 3 – figure supplement 2-source data 1* file. **Figure 3 – figure supplement 2-source data 1**: relates to panels A-C.

**Figure 3 – figure supplement 3.**
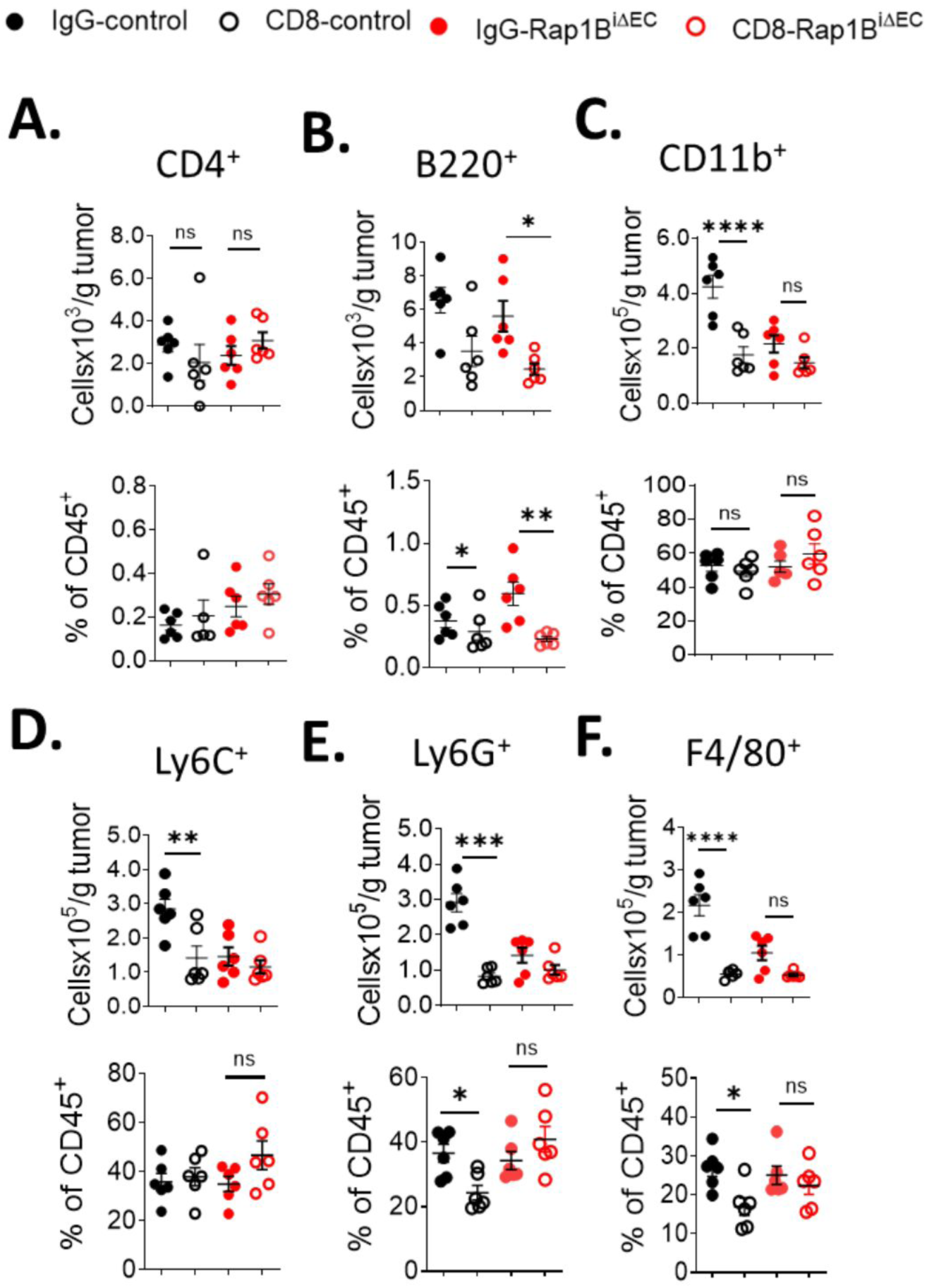
Effect of CD8^+^ T-cell depletion on tumor-infiltra:ng lymphocytes (TIL) in control and Rap1B^iΔEC^ mice. Immunophenotypic examination showing quantitation of **(A)** CD4^+^T cells, **(B)** B cells (B220^+^), **(C)** myeloid cells (CD11b^+^), **(D)** monocytes (Ly6C^+^), **(E)** neutrophils (Ly6G^+^) and **(F)** macrophage (F4/80^+^) in Rap1B^iΔEC^/control mice treated with either IgG isotope or CD8 mAb. Data are presented as the mean ± S.E.M. *, P < 0.05, **, P < 0.01. n = 6 mice per group, one-way ANOVA with Tukey’s multiple comparisons post hoc test. Data and statistical outputs are available in *Figure 3 – figure supplement 3-source data 1* file. **Figure 3 – figure supplement 3-source data 1**: relates to panels A-F.

**Figure 3 – figure supplement 4.**
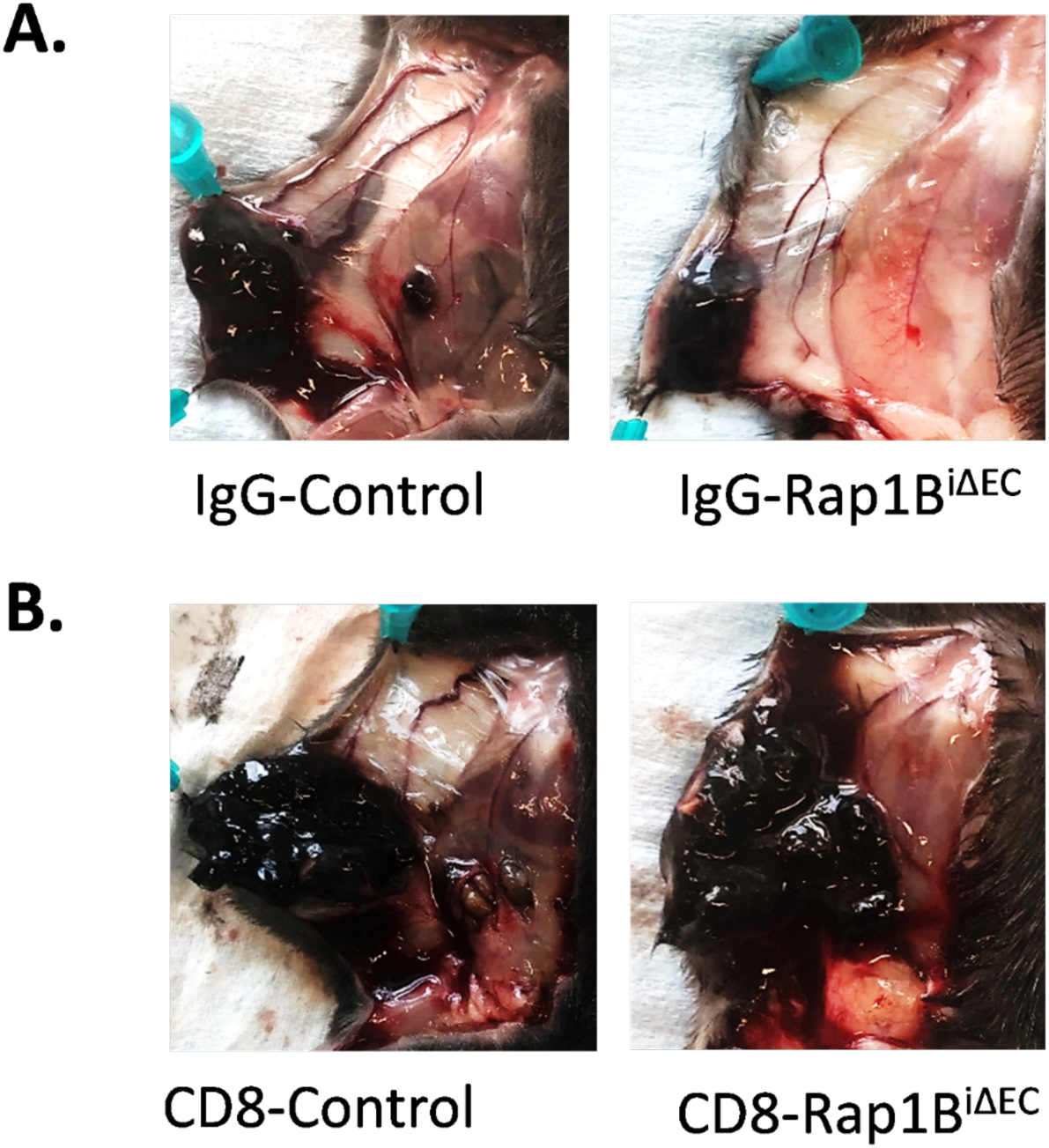
Depletion of CD8^+^ T-cells normalizes tumor growth in Rap1B^iΔEC^ mice. Representative photographs showing pattern of tumor growth in Rap1B^iΔEC^/control mice treated with either IgG isotope or CD8 mAb. **(A)** IgG isotope has no effect on tumor growth in control group. **(B)** CD8^+^ T cell depletion significantly rescued tumor reduction in Rap1B^iΔEC^ mice group.

**Figure 4 – figure supplement 1.**
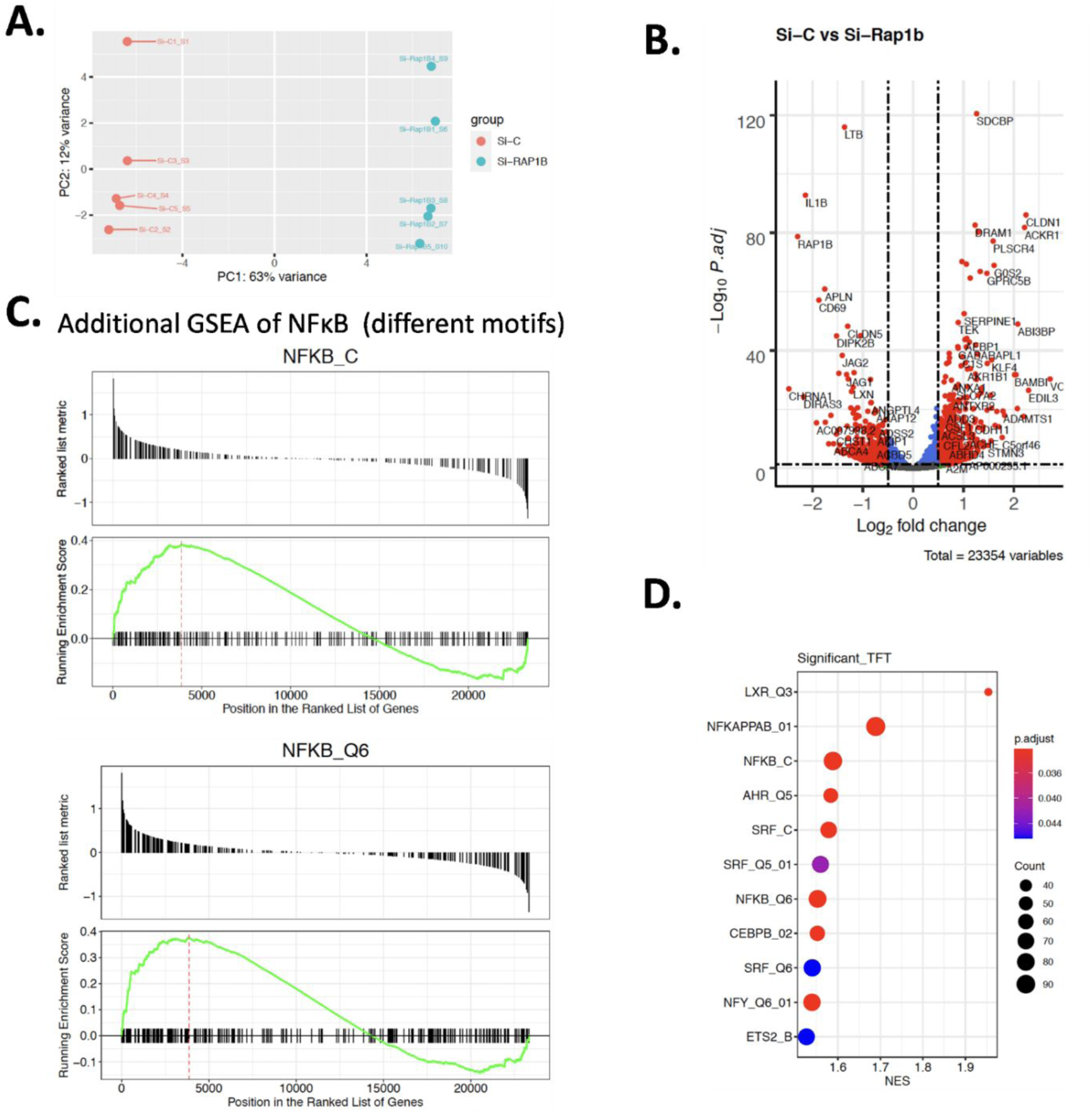
Transcriptomic analysis of TNF-α-s:mulated siControl and siRap1B ECs. **(A)** Principal component analysis of samples. (**B**) Volcano plot of differential gene expression, the top 50 significantly differentially expressed genes are labeled. (**C**) Gene set enrichment analyses (GSEA) of NFκB target genes. (**D**) Dot plot of top significantly enriched transcription factors by GSEA (n = 5). Link to sequence data deposited within NCBI GEO is available in *Figure 4-source data 1* file.

**Figure 5 – figure supplement 1.**
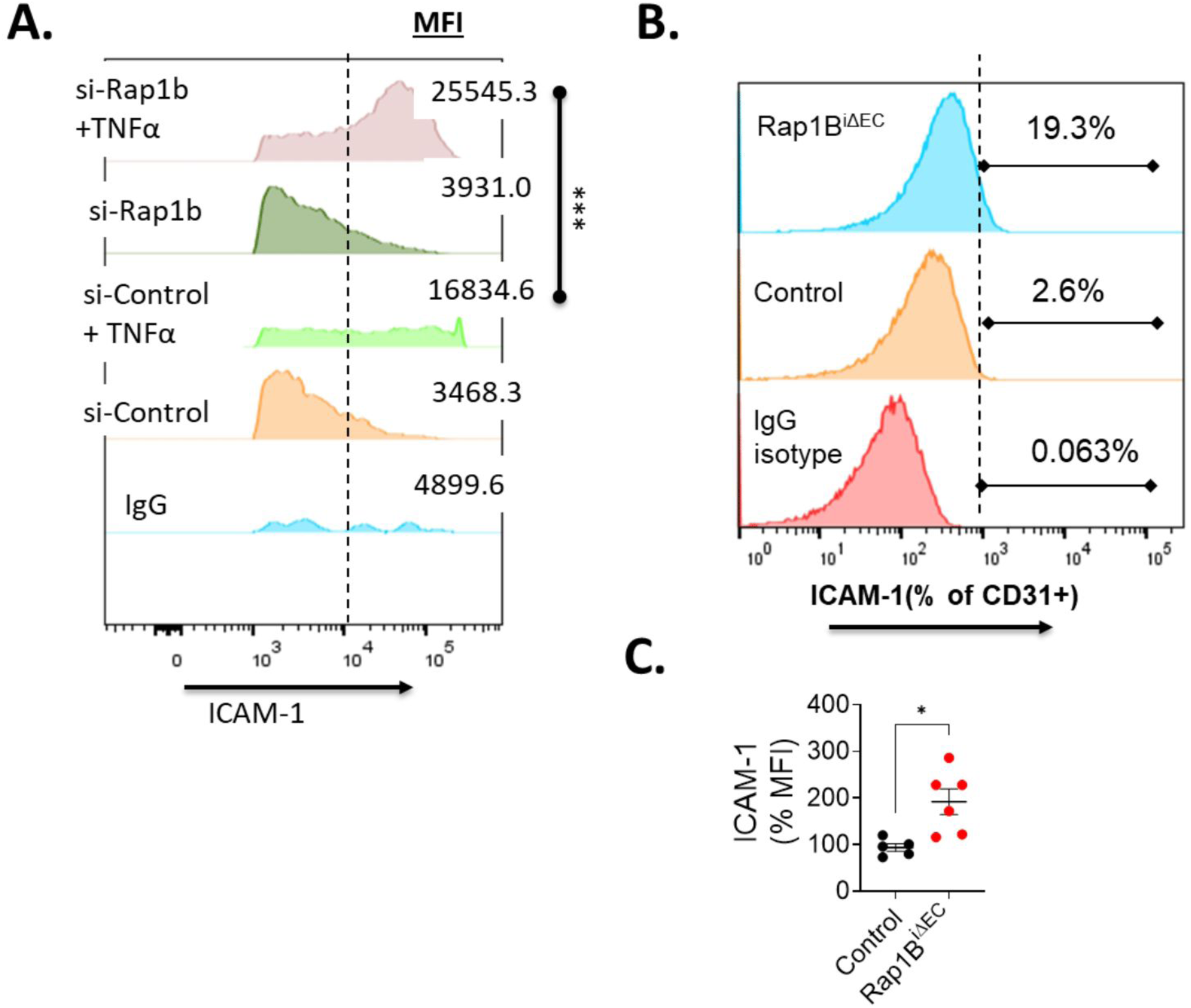
Elevated ICAM-1 surface expression in Rap1B-deficient tumor ECs and HUVECs. **(A)** Flow cytometric analysis of ICAM-1 expression on siRap1B and siControl HUVEC cells. (**B**) Flow cytometric analysis of ICAM-1 expression on tumor ECs from Rap1B^iΔEC^ and control mice: representative flow cytometry histogram and median fluorescence intensity (MFI) plot (**C**). Data are presented as the mean ± S.E.M. **, P < 0.01, Student’s t test. Data and statistical outputs are available in *Figure 5 – figure supplement 1-source data 1* file. **Figure 5 – figure supplement 1-source data 1:** relates to panels A and C.

---

**Supplemental Table 1:** Key Resources and Antibodies.

| Reagent type (species) or resource | Designation | Source or reference | Identifiers | Additional information |
| --- | --- | --- | --- | --- |
| (Mus musculus) | VE-Cadh-CreN-Rap1a<sup>+/+</sup>Rap1b<sup>f/f</sup> | In house breeding |  | C57Bl/6 genetic background |
| (Mus musculus) | VE-Cadh-CreY-Rap1a<sup>+/+</sup>Rap1b<sup>f/f</sup> | In house breeding |  | C57Bl/6 genetic background |
| (Mus musculus) | Rap1B<sup>(+/-)</sup> | In house breeding |  | C57Bl/6 genetic background |
| (Mus musculus) | Rap1B<sup>(+/+)</sup> | In house breeding |  | C57Bl/6 genetic background |
| Other | HUVECs( Human Umbilical Vein ECs) | ATCC | Cat# CRL-1730<br>RRID:CVCL_2959 |  |
| Other | B16F10 | ATCC | Cat# CRL-6475<br>RRID:CVCL_0159 | Mouse melanoma cell line |
| Other | Jurkat E6–1 | ATCC | Cat# TIB-152 | Human T cell leukemia cell line |
| Other | Rap1B siRNA | Dharmacon |  | 50nM in reaction |
| Other | Control siRNA | Dharmacon |  | 50nM in reaction |
| Chemical compound, drug | TNF-α | Sigma-Aldrich | Cat# H8916-10UG | 50nM in assay |
| Chemical compound, drug | VEGF | Sigma-Aldrich | Cat# V7259 | 50 ng/ml in assay |
| Chemical compound, drug | Calcein Blue AM | BD Biosciences | Cat# 564060 | 5uM in assay |
| Other | pNL3.2.NF-κB-RE[NlucP/NF-κB-RE/Hygro] Vector | Promega | Cat#N1111 | 2 ug in assay |
| Other | pNL3.2.CMV | Promega | Cat#N1411 | 2 ug in assay |
| Assay kit | Nano-Glo® Luciferase Assay System | Promega | Cat.# N1110 |  |
| Antibody | VCAM-1 (clone E1E8X) Rabbit mAb antibody | Cell Signaling Technology | Cat# 13662,<br>RRID:AB_2798286 | WB (1:2000) |
| Antibody | Rap1A/Rap1B (clone 26B4) Rabbit mAb antibody | Cell Signaling Technology | Cat# 2399,<br>RRID:AB_2284915 | WB (1:2000) |
| Antibody | β-Actin mouse, rat, human (clone C4) mAb antibody | Santa Cruz Biotechnology | Cat# sc-47778<br>RRID:AB_2714189 | WB (1:500) |
| Software, algorithm | GraphPad Prism version 5.02 | GraphPad Software |  |  |
| Software, algorithm | Flow Jo software |  |  |  |
| Antibody | FITC anti-mouse CD3, clone: 145-2C11 | Bio Legend | Cat# 100305,<br>RRID:AB_312670 | Dilution: 1:300<br>For Tumor Infiltrate Lymphocyte (TIL) |
|  | (Armenian Hamster-monoclonal) |  |  | analysis by flow cytometry |
| Antibody | APC anti-mouse CD4, Clone GK1.5, (rat-monoclonal) | Bio Legend | Cat# 100412, RRID:AB_312697 | Dilution: 1:400<br>For TIL analysis by flow cytometry |
| Antibody | FITC anti-mouse CD4, Clone GK1.5, (rat-monoclonal) | Bio Legend | Cat# 100405, RRID:AB_312690 | Dilution: 1:400<br>For TIL analysis by flow cytometry |
| Antibody | Brilliant Violet 510(TM) anti-mouse, Clone M1/70, (rat-monoclonal) | Bio Legend | Cat# 101263, RRID:AB_2629529 | Dilution: 1:400<br>For TIL analysis by flow cytometry |
| Antibody | PE/Cyanine7 anti-mouse CD31, Clone 390, (rat-monoclonal) | Bio Legend | Cat# 102418, RRID:AB_830757 | Dilution: 1:300<br>For TIL analysis by flow cytometry |
| Antibody | APC/Cyanine7 anti-mouse CD44, Clone IM7, (rat-monoclonal) | Bio Legend | Cat# 103028, RRID:AB_830785 | Dilution: 1:400<br>For TIL analysis by flow cytometry |
| Antibody | PE/Cyanine5 anti-mouse CD45, Clone 30-F11, (rat-monoclonal) | Bio Legend | Cat# 103109, RRID:AB_312974 | Dilution: 1:400<br>For TIL analysis by flow cytometry |
| Antibody | APC/Cyanine7 anti-mouse CD45, Clone 30-F11, (rat-monoclonal) | Bio Legend | Cat# 103116, RRID:AB_312981 | Dilution: 1:400<br>For TIL analysis by flow cytometry |
| Antibody | PE/Dazzle(TM) 594 anti-mouse, clone RA3-6B2, (rat-monoclonal) | Bio Legend | Cat# 103258, RRID:AB_2564053 | Dilution: 1:200<br>For TIL analysis by flow cytometry |
| Antibody | Pacific Blue(TM) anti-mouse F4/80, Clone BM8, (rat-monoclonal) | Bio Legend | Cat# 123124, RRID:AB_893475 | Dilution: 1:300<br>For TIL analysis by flow cytometry |
| Antibody | Alexa Fluor(R) 700 anti-mouse Ly-6G, Clone 1A8, (rat-monoclonal) | Bio Legend | Cat# 127622, RRID:AB_10643269 | Dilution: 1:300<br>For TIL analysis by flow cytometry |
| Antibody | PE anti-mouse Ly-6C, Clone HK1.4, (rat-monoclonal) | Bio Legend | Cat# 128007, RRID:AB_1186133 | Dilution: 1:300<br>For TIL analysis by flow cytometry |
| Antibody | PerCP anti-mouse Ly-6G/Ly-6C (Gr-1), Clone RB6-8C5, (rat-monoclonal) | Bio Legend | Cat# 108425, RRID:AB_893560 | Dilution: 1:400<br>For TIL analysis by flow cytometry |
| Antibody | APC anti-mouse NK-1.1, Clone PK136, (rat- monoclonal) | Bio Legend | Cat# 108709, RRID:AB_313396 | Dilution: 1:200<br>For TIL analysis by flow cytometry |
| Antibody | PE anti-mouse CD8a, Clone 53-6.7, (rat-monoclonal) | Thermo Fisher Scientific | Cat# 12-0081-82, RRID:AB_465530 | Dilution: 1:200<br>For TIL analysis by flow cytometry |
| Antibody | eFluor 450 anti-mouse CD106 (VCAM-1), Clone 429, (rat-monoclonal) | Thermo Fisher Scientific | Cat# 48-1061-82, RRID:AB_1633378) | Dilution: 1:300<br>For TIL analysis by flow cytometry |
| Antibody | FITC anti-mouse CD54 (ICAM-1), Clone YN1/1.7.4, (rat-monoclonal) | Thermo Fisher Scientific | Cat# 11-0541-82, RRID:AB_465094) | Dilution: 1:300<br>For TIL analysis by flow cytometry |
| Antibody | PE cy7 anti-mouse Granzyme B, Clone NGZB, (rat-monoclonal) | Thermo Fisher Scientific | Cat# 25-8898-82, RRID:AB_10853339 | Dilution: 1:200<br>For TIL analysis by flow cytometry |
| Antibody | eFluor 450-Rat IgG1 κ Isotype control, Clone eBRG1, (rat-monoclonal) | Thermo Fisher Scientific | Cat# 48-4301-82, RRID:AB_1271984 | Dilution: 1:200<br>For FMO control in flow cytometry |
| Antibody | PE cy7-Rat IgG2a κ Isotype control, Clone eBR2a, (rat-monoclonal) | Thermo Fisher Scientific | Cat# 25-4321-82, RRID:AB_470200 | Dilution: 1:200<br>For FMO control in flow cytometry |
| Antibody | FITC-Rat IgG2b κ Isotype control, Clone eB149/10H5, (rat-monoclonal) | Thermo Fisher Scientific | Cat# 11-4031-82, RRID:AB_470004) | Dilution: 1:200<br>For FMO control in flow cytometry |

